# CellCover Defines Marker Gene Panels Capturing Developmental Progression in Neocortical Neural Stem Cell Identity

**DOI:** 10.1101/2023.04.06.535943

**Authors:** Lanlan Ji, An Wang, Shreyash Sonthalia, Seungmae Seo, Daniel Q. Naiman, Laurent Younes, Carlo Colantuoni, Donald Geman

## Abstract

1

Definition of cell classes across the tissues of living organisms is central in the analysis of growing atlases of single-cell RNA sequencing (scRNA-seq) data across biomedicine. Marker genes for cell classes are most often defined by differential expression (DE) methods that serially assess individual genes across landscapes of diverse cells. This serial approach has been extremely useful, but is limited because it ignores possible redundancy or complementarity across genes that can only be captured by analyzing multiple genes simultaneously. Interrogating binarized expression data, we aim to identify discriminating *panels* of genes that are specific to, not only enriched in, individual cell types. To efficiently explore the vast space of possible marker panels, leverage the large number of cells often sequenced, and overcome zero-inflation in scRNA-seq data, we propose viewing marker gene panel selection as a variation of the “minimal set-covering problem” in combinatorial optimization. Using scRNA-seq data from blood and brain tissue, we show that this new method, CellCover, performs as good or better than DE and other methods in defining cell-type discriminating gene panels, while reducing gene redundancy and capturing cell-class-specific signals that are distinct from those defined by DE methods. Transfer learning experiments across mouse, primate, and human data demonstrate that CellCover identifies markers of conserved cell classes in neocortical neurogenesis, as well as developmental progression in both progenitors and neurons. Exploring markers of human outer radial glia (oRG, or basal RG) across mammals, we show that transcriptomic elements of this key cell type in the expansion of the human cortex likely appeared in gliogenic precursors of the rodent before the full program emerged in neurogenic cells of the primate lineage. We have assembled the public datasets we use in this report within the NeMO Analytics multi-omic data exploration environment [1], where the expression of individual genes ([NeMO: Individual genes in cortex](https://nemoanalytics.org/p?&l=CellCover&g=HOPX) and [NeMO: Individual genes in blood](https://nemoanalytics.org/p?&l=immune&g=CSF1R)) and marker gene panels ([NeMO: Telley 3 CellCover Panels](https://nemoanalytics.org/p?p=p&l=CellCover&c=TelleyCellCover3sets&algo=binary), [NeMO: Telley 12 CellCover Panels](https://nemoanalytics.org/p?p=p&l=CellCover&c=TelleyCellCover12sets&algo=binary), [NeMO: Sorted Brain Cell CellCover Panels](https://nemoanalytics.org/p?p=p&l=CellCover&c=LiuCellCoverSorted&algo=binary), and [NeMO: Blood 34 CellCover Panels](https://nemoanalytics.org/p?p=p&l=immune&c=CellCoverHao34&algo=binary)) can be freely explored without coding expertise. CellCover is available in [CellCover R](https://github.com/lanlanji/CoveringPackage) and [CellCover Python](https://pypi.org/project/CellCover/).

**Graphical Abstract:** 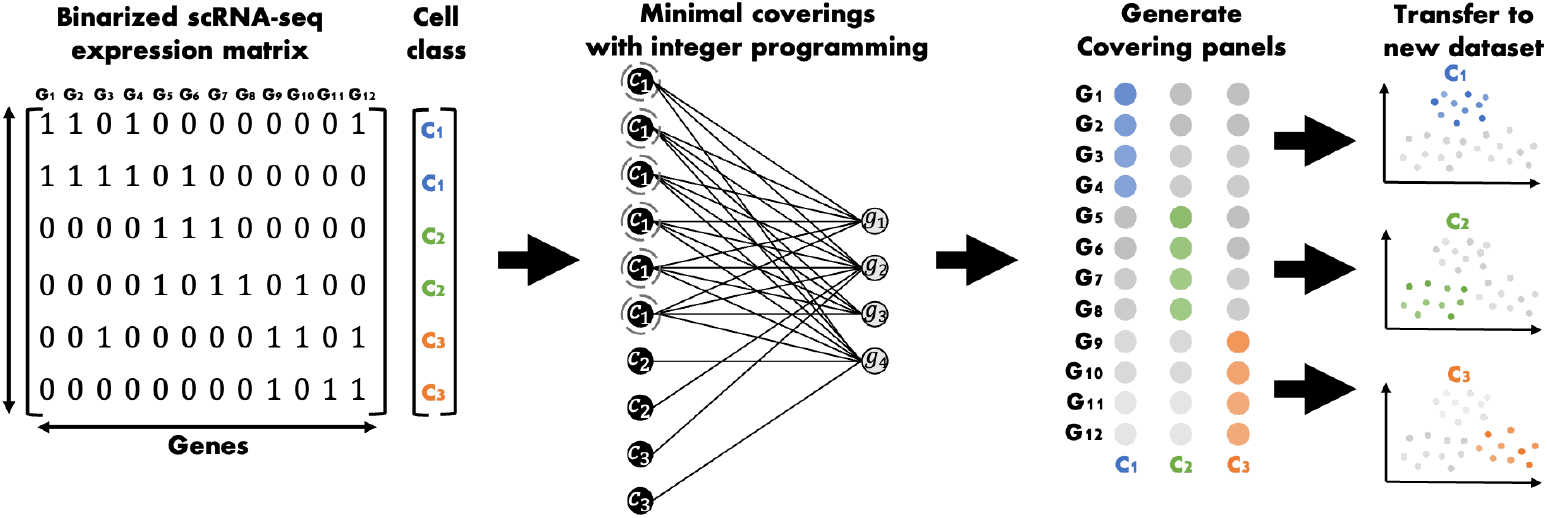

## 2 Introduction

scRNA-seq technologies provide measurements of RNA molecules in many thousands of individual cells, yielding a rich source of information for determining attributes of cell populations, such as cell types and the variation in gene expression from cell to cell, which are not available from bulk RNA sequencing data [2, 3, 4, 5, 6]. After standard preprocessing and normalization, a core challenge in the analysis of scRNA-seq data is to annotate the cells in a given dataset with a label indicating cell class, type or state. This usually involves multiple phases of processing, notably dimension reduction, clustering, and finding “marker genes” for the labels of interest. Markers are usually selected by first ranking individual genes based on a statistical test of differential expression (DE) across cell type labels and then selecting a panel of top-ranking genes based on manual curation, estimates of predictive capacity, a priori information (e.g., cell-surface proteins), or simply panel size [7, 8, 9, 10, 11, 12, 6, 13]. Importantly, the construction of such marker panels incorporates data from only one gene at a time, i.e., the panel is assembled “gene by gene,” and does not take into account possible gene combinations that could better distinguish cell classes.

Complications arise because scRNA-seq data are high dimensional [14], highly heterogeneous, generally noisier than bulk RNA-seq, and stochastically zero-inflated. Complex pipelines with many choices require manual intervention [7, 15], and raise issues of bias and reproducibility [16, 17]. In order for downstream analyses to be computationally feasible, restricting dimension, such as using small panels of marker genes, is necessary. Analysis of scRNA-seq data is generally done at the univariate (single gene) level, even though the relevant biology is often decidedly multivariate. In particular, probability distributions, when proposed, are for the expression of individual genes (i.e., marginal distributions) not for gene panels (i.e., higher-order marginals). These analyses do not take into account how marker genes may cooperate to define cell types, or possible heterogeneity within cell types.

To avoid these limitations, we formulate marker gene selection as a variation of the well-known “minimal set-covering problem” in combinatorial optimization. In our case, the “covering” elements are genes, and the objects to be covered are the cells in a particular class. A cell is covered at depth ***d*** by a gene panel if at least ***d*** genes in this panel are expressed (raw count ≥ 1) in the cell (see details in Supplemental section 1). A set of genes is then a depth ***d*** marker panel for a cell type if nearly all cells of that type are covered at depth ***d***. We define the covering rate as 1 − ***α***, where ***α*** represents the fraction of uncovered cells, i.e. the false negative rate. This covering rate can be adjusted to balance specificity and sensitivity of the marker genes. Prior to selection of the gene panel, a weight is calculated for each gene, defined as its expression outside the target class divided by its expression within that class. Lower weights, therefore, correspond to greater discriminative power. This weight calculation can be carried out as one class versus all other cells (as used in this report), or as one class versus its closest neighboring class, and can be based on either binary or count-based expression. The optimal marker panel is then selected from genes with higher expression within the cell type of interest (weight < 1), determined by minimizing the sum of weights in the gene panel while ensuring that the panel achieves the desired covering rate. We formulate this search as a constrained optimization problem constructed as an integer linear program ([18], [19]; see Methods section 5) This approach enables the identification of compact sets of marker genes that reliably distinguish cell populations. Additionally, gene panels can be transferred across studies to explore shared cell-type-specific expression patterns by examining covering depth of the panels in new cells and covering rates in new cell classes.

The minimal covering approach in CellCover is designed specifically to leverage the high number of cells routinely observed in scRNA-seq data in order to be robust to the stochastic zero-inflation known to be pervasive in this data modality: Because the covering algorithm optimizes a set of genes rather than individual markers and different true marker genes will have drop-outs in different cells, it follows that if many cells of a class are sequenced, zero-inflation can be overcome (while also mitigating the impact of higher noise at low expression levels). In contrast, standard methods, based on selecting genes for which a differential expression statistic is extreme, cannot borrow strength across genes in a marker panel when selected gene-by-gene. CellCover searches for small panels of covering marker genes that precisely define cell classes together as a set, and are also expandable, i.e., can be readily linked to additional related genes for the interrogation of cell-type-specific function. In addition, CellCover output can be used in gene set over-representation analysis and the deeper exploration of heterogeneity within individual cell classes.

To demonstrate CellCover’s utility in identifying cell-type-specific markers, we benchmark CellCover against leading marker selection methods using single-cell RNA-seq datasets from human blood. We show that CellCover effectively reduces marker redundancy and outperforms most methods in predicting cell types. In a series of *in silico* experiments we use CellCover to explore cell classes and their developmental progression across neurogenesis in the excitatory neuronal lineage of the mammalian neocortex. Through cross-dataset marker transfer experiments, we validate the robustness and transferability of CellCover markers across various sequencing protocols and biological contexts, highlighting its capacity to consistently map cell-type-specific gene expression from a reference dataset to cells or samples in new target datasets.

## 3 Results

### 3.1 CellCover on blood scRNA-seq data: Benchmark and Transfer

#### 3.1.1 Benchmarking cell type definition

We compare CellCover with leading marker gene selection algorithms - Wilcoxon rank sum based differential expression (DE) [20], RankCorr [21], scGeneFit [15], and PhenotypeCover [22] - to evaluate their effectiveness in identifying cell state resolving markers. This benchmark analysis uses the CBMC CITE-Seq dataset [23], which provides a rich source of immune cell information from cord blood mononuclear cells (CBMCs) through multimodal single-cell analysis. To systematically assess the different methods, we employ a five-fold cross-validation approach to first extract marker genes from the training data, using only the RNA expression, and then test cell type label recovery in test data using these markers in a support vector machine (SVM). To generate marker gene panels, we implemented CellCover at various depths of coverage with a cover rate 90% (***α*** = 0.1), using both log-normalized and binary expression data to define gene weights (see details in Method section 5.3.3). For DE, we selected the top genes for each cell type, matching the exact number of markers in the CellCover panel for that cell type. Unlike DE and CellCover, the RankCorr, scGeneFit, and PhenotypeCover methods do not generate cell-type-specific marker panels but rather global panels that distinguish all cell types. Therefore, at each covering depth, we select an equivalent number of markers for RankCorr, scGeneFit, and PhenotypeCover to match the global CellCover panel size (i.e. the total number of unique marker genes used for all cell classes combined). After selecting marker gene panels for all methods we trained an SVM on the training data using only the marker genes identified by each method, and then report the average balanced accuracy on the test data restricted to these markers (see details in Method section 5.3). The results in Figure 1A show that CellCover, along with DE methods, consistently achieves the highest balanced accuracy at different marker panel sizes. Importantly, while CellCover and DE methods achieve similar levels of performance in cell type definition, they do so by utilizing distinct sets of genes, selecting only 20-30% of the same genes in their marker panels (Figure 1B, see detail in Method Section 5.3.4). This indicates that CellCover and DE methods leverage quite different elements of gene expression to distinguish cell types.

**Figure 1:**
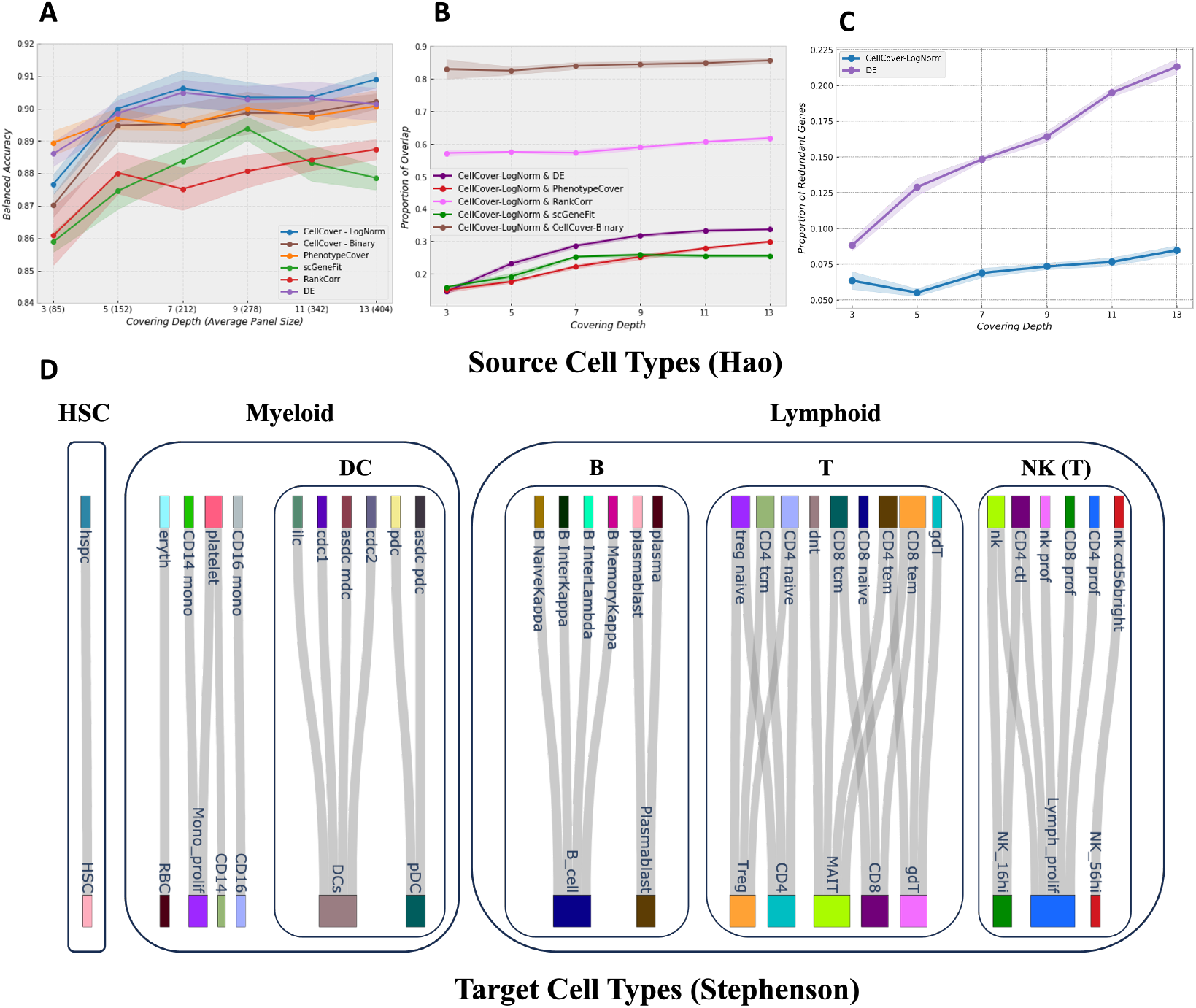
Benchmarking performance of CellCover and competing methods on the CBMC dataset & Blood Cell Type Mapping. A) Balanced accuracy versus the size of the marker panel for all methods, with standard deviation across random seeds shown as shaded areas. CellCover implemented with weights calculated from log normalized or binary data are both shown. B) Proportion of intersection between each method’s global marker panel and the CellCover (log-normalized) reference panel C) Proportion of redundant marker genes selected for multiple cell classes versus panel size for CellCover and DE, also including standard deviation across random seeds. D) Sankey diagram of the mapping from source blood cell types [24] to target blood cell types [25]. Original authors’ cell type labels are used: HSC = hematopoietic stem cells, DC = dendritic cells, B = B-cells, T = T-cells, NK = natural killer cells, eryth + RBC = red blood cells, mono= monocytes, ilc = innate lymphoid cells, prof + prolif = proliferating.

In addition to optimal cell class label recovery, it is desirable that marker gene panels do not contain the same genes for different cell classes. Because DE methods choose marker genes one-at-a-time from a ranked list of statistics for each cell class, they often select the same gene for more than one cell class. This is especially if some cell classes are more similar to one another than to other classes, which is a common scenario in many biological contexts. As evidenced by Figure S2, CellCover selects fewer redundant marker genes compared to DE at all covering depths, indicating that its marker panels capture more unique, non-overlapping signals across cell types. Figure 1C provides another view of this, showing that the proportion of redundant CellCover markers remains stable around 7.5%, while the proportion of redundant DE markers steadily increases with the marker panel size, surpassing 20% in larger panels (see detail in Method Section 5.3.5). This greatly reduced redundancy demonstrates that CellCover generates concise marker panels in which genes provide distinct and complementary information.

#### 3.1.2 Cell type mapping across datasets through marker transfer

To demonstrate that CellCover effectively captures robust cell-type-specific information that is transferable across diverse single-cell RNA-seq datasets, we performed cross-dataset marker transfer experiments between datasets derived from human blood. Using the Hao PBMC CITE-seq dataset [24] as a source dataset with marker panels generated at depth 5 and a covering rate of 98%, we evaluated the ability of these CellCover marker panels to correctly identify cell types in four new target datasets: The Tabula Sapiens blood data [26] sequenced with three protocols (smart-seq2, 10x 3’, and 10x 5’) and the Stephenson COVID-19 PBMC dataset [25], restricted to cells from healthy controls (see Supplemental section 9; [NeMO: Individual genes in blood](https://nemoanalytics.org/p?l=immune&g=CSF1R) and [NeMO: Blood 34 CellCover Panels](https://nemoanalytics.org/p?p=p&l=immune&c=CellCoverHao34&algo=binary)).

The Sankey diagrams in Figure 1D and Figure S15-17 (see details in Method section 5.4) demonstrate that marker gene panels identified by CellCover accurately map cell types from the source dataset to cell types of the same hematopoetic lineage in the target datasets (despite divergent cell type labeling of the original studies). This can be seen in Figure 1D where cell types of common lineage origin are very rarely incorrectly mapped i.e. cell type mappings stay within hematopoetic lineages noted in the figure. The one exception to this is that a small proportion of the dendritic cells (DCs) in the target dataset were mapped to innate lymphoid cells (ilc), a cell type that is not well-defined and extremely rare in the circulating blood [27]. This result indicates the robustness of CellCover marker gene panels in mapping cell-type-specific signals across different sequencing technologies and experiments.

### 3.2 scRNA-seq datasets in neocortical neurogenesis

We illustrate the use of CellCover in the context of dorsal telencephalic neurogenesis during mid-gestation, when the majority of excitatory neurons of the neocortex are produced. The mature neocortex in mammals is made up of 6 layers of post-mitotic neurons. Neurons of each new layer are produced in succession, with neurons of the new layer migrating past previously created layers, leading to an “inside-out” arrangement of neurons by birthdate [28], with deep layers (VI-V) being born first and superficial layers (II-IV) born last. Neurons are generated sequentially from radial glia (RG), the neural stem cells of the telencephalon, and then intermediate progenitor cells (IPCs) through both symmetric and asymmetric cell division [29]. The number and diversity of neural stem and progenitor cells in the cortex have been greatly expanded in the evolution of the primate lineage. In particular, the appearance of the outer subventricular zone (OSVZ) in primates appears to underly the development of the supragranular layers of the cortex (layers II and III) which contribute to higher cognition through extensive cortico-cortico connections [30]. Notably, this includes greater proliferative capacity of neural stem cells and the expansion of outer radial glia (oRG, or basal radial glia, bRG), which are rare in rodents but numerous in primates and especially in humans [31, 32]. As each cortical layer is generated, newborn neurons arise contemporaneously from progenitor cells in the germinal zones (GZ) and migrate radially along processes of the RG, outward toward the surface of the telencephalon, to the cortical plate (CP), coming to rest together at their characteristic position in the developing cortex [33]. Neurons arising at the same time and place, and therefore from precursors with common molecular features, share unique morphological and functional characteristics. We are interested both in the general process by which neural precursor cells yield post-mitotic neurons, and in the more nuanced dynamics by which specific properties are imparted to neurons which are born together and which come to rest in the same cellular microenvironment, i.e., cortical region and layer.

How the molecular dynamics of neural progenitor identity across this progression impact the fate and function of the neurons being produced is not fully understood. To explore both the principal cell identities and the developmental progression within cell types, we first focus on scRNA-seq data from Telley et al. [34] containing cells extracted during neocortical neurogenesis in the mouse, and for which highly precise developmental metadata are available. This dataset contains expression data from cells produced across four days of mid-gestational embryonic development during the peak of excitatory neocortical neurogenesis (E12-15). Using a technique to specifically label ventricular radial glia (RG) cells during their final neurogenic cell division, cells born on each of the four embryonic days were sampled at one hour, 24 hours, and 96 hours after this terminal division. This yielded a two-dimensional temporal indexing of the data based on the embryonic age of the animals and the precise time since the final division of the individual cells sequenced. We define CellCover marker gene panels in the principal cell types involved in neocortical neurogenesis as they progress through development and then validate their ability to capture conserved cell-type-specific signals in additional scRNA-seq data from developing mouse [35, 36], primate [37, 38], and human [39, 40] cortical tissue. In order to focus on cell states and transitions in excitatory neurogenesis, for all the studies we use in this report, we attempt to analyze only cells of the excitatory neurogenic lineage, omitting inhibitory neurons and later astroglial and oligodendroglial cells, as well as non-neural cells including microglia and endothelial cells. Individual gene expression data in all the public studies we employ here can be explored without coding expertise at: NeMO: Individual genes in cortex.

### 3.3 CellCover marker gene panels in mouse neocortical development #1: Cell types

The precise cell birth dating that is coupled to scRNA-seq data in the Telley study [34] provides a unique opportunity to interrogate cell type transitions during neocortical neurogenesis. To take advantage of this experimental design, rather than using data-driven clusters or biologist-annotated cell labels, we employed the empirical timing data for individual cells to define cell classes. At 1 hour after final division, it is likely that the transcriptome of RGs is still representative of the progenitor state, while at 96 hours, we expect cells to have transitioned to a postmitotic neuronal state. Cells at 24 hours after final division represent transitional states between the RGs and postmitotic neurons, likely spanning intermediate progenitor, neuroblast and nascent neuronal states. Using our novel CellCover approach, we generated marker gene panels defining cell states at 1, 24, and 96 hours after the terminal division of ventricular RGs (regardless of embryonic day).

Figure 2A depicts the gene panels derived from cells at each of the three times sampled, shown as the proportion of cells at each time that express each of the marker genes at non-zero levels. The original authors noted elevated heterogeneity in the 24H cell class, which can be seen as more frequent expression of 24H cell markers in the other time classes. This is consistent with the 24H cells having their own transcriptomic characteristics, while also harboring elements of both the preceding progenitor state and the subsequent neuronal state. The expression of these maker gene panels can be viewed across the public datasets we use in this report at: [NeMO: Telley 3 CellCover Panels](https://nemoanalytics.org/p?p=p&l=CellCover&c=TelleyCellCover3sets&algo=binary).

**Figure 2:**
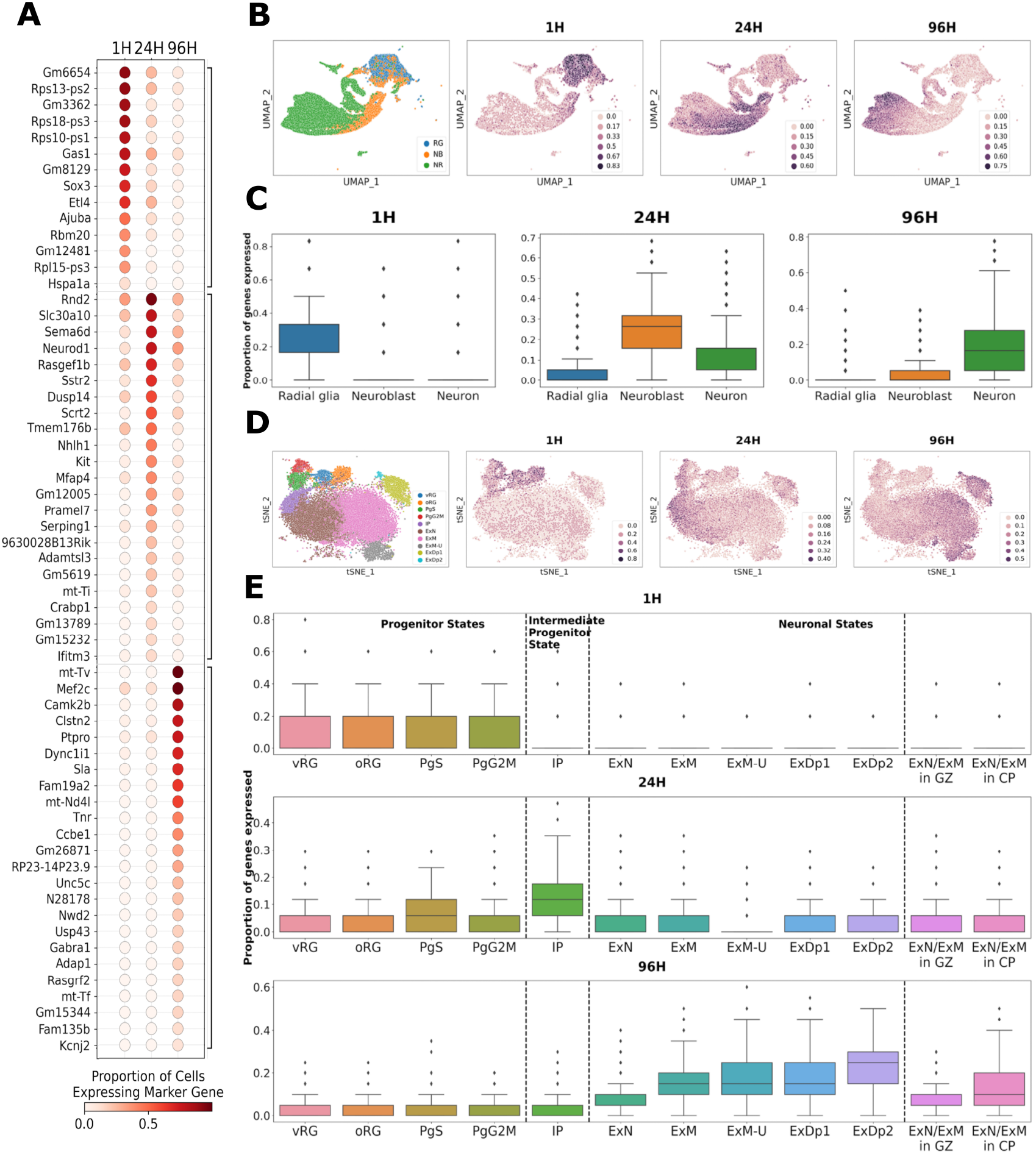
CellCover marker gene panels in the developing mammalian neocortex. **A**. Dot plot of empirical conditional expression probabilities of CellCover marker panels of each cell age. The marker genes of each cell age are grouped along the horizontal axis and sorted by expression frequency in the cell class of interest. The color of the dots represents the expression probability of markers conditioned on time since a cell’s terminal division, i.e., the proportion of cells within a class expressing the marker gene. For this analysis, cells of each time point were pooled across embryonic ages (E12–15). The panel is obtained using the CellCover with ***α*** = 0.02 and ***d*** = 5. RG = radial glia, NB = neuroblast, NR = neuron. **B**. Transfer of marker gene panels from the Telley data [34] to a second mouse neocortex dataset shows consistent identification of the primary cell types in mouse neurogenesis. Left: UMAP representations of cells from the dorsal forebrain excitatory lineage in the LaManno atlas of mouse brain development [35], colored by cell labels assigned by the original authors. This is followed by the same UMAPs, now showing the proportion of gene panels derived from the Telley dataset, labeled 1H, 24H and 96H, expressed at non-zero levels in each individual cell. These last three plots illustrate the transfer of marker gene panels derived from the Telley data to the LaManno data. **C**. Box plots of these same proportions broken down by cell-type labels provided by the original authors. **D**. Transfer of marker gene panels from the Telley data in mouse to data in the developing human neocortex from Polioudakis et al. [39], shows the identification of conserved cell types in neurogenesis across mammalian species. Left: tSNE map of cells from the dorsal forebrain excitatory lineage in the Polioudakis data. This is followed by the same tSNE maps, now showing the transfer of marker gene panels derived from progenitors labeled 1, 24, and 96 hours after terminal cell division in the Telley data. The map is colored by the proportion of each gene panel that is expressed at non-zero levels in each individual cell of the human Polioudakis dataset. Original athor cell labels are used: RG = radial glia, v = ventricular, o = outer, Pg = cycling progenitor, S = in S phase, G2M = in G2M phase, IP = intermediate progenitor, Ex = excitatory neuron, N = new migrating, M = maturing, Dp = deep layer, U = upper layer.**E**. Box plots of these same proportions broken down by cell type labels and microdissection information provided by the original authors. The final two boxes indicate expression in neuronal subtypes segregated by physical location: germinal zone (GZ) or cortical plate (CP) microdissection.

These marker gene panels contain canonical markers of neural progenitors, intermediate progenitors / neuroblasts and neurons, including Sox3, Neurod1, and Mef2c in 1H, 24H, and 96H cells, respectively. The panels also contain many lesser known genes (Gm …), pseudogenes (… -ps), and mitochondrialy encoded genes (mt-…) which are expressed at much lower levels, but together precisely distinguish these neurogenic cell classes. Mitochondrial transcripts, and tRNAs in particular, are often disregarded as uninformative “housekeeping” genes or even indications of low-quality RNA samples. These are dangerous assumptions, especially when exploring neurogenesis where the mitochondrial content of cells changes dramatically as cells transition from stem and progenitor states to post-mitotic neurons. The expression of mt-Tv, for example, is highly specific to 96H cells (Figure 2A), and while often disregarded as a housekeeping tRNA, it also forms a structural component of the mitochondrial ribosome [41] and is transcribed in higher abundance as energy demands of nascent neurons increase. This suggests that many genes not often selected by conventional marker gene finding methods (and often intentionally filtered out by biologists) hold significant cell-type-specific information. To assess this possibility and the utility of CellCover marker gene panels in general, we next examine the ability of these panels to interrogate cell-type-specific signals in additional neocortical datasets.

### 3.4 Transfer of CellCover marker gene panels #1: Conserved cell types in neocortical neurogenesis

To assess the ability of CellCover marker gene panels to capture consistent cell-type-specific signals in murine neurogenesis, we explored additional *in vivo* data from the dorsal mouse telencephalon. We examined the expression of CellCover panels derived from the Telley mouse dataset in a recent atlas of the developing mouse brain (La Manno et al. [35]; Figure 2B and 2C). For both these mouse scRNA-seq datasets the precise gestational age of the mouse pup of origin is known for all cells collected. However, while there are ground truth labels of the time from ventricular progenitor terminal cell division in the Telley data, in the atlas we must rely on cell type annotation provided by the researchers, which include radial glia, neuroblast, and neuron classes sequentially along the neurogenic trajectory. As expected, marker genes derived from progenitors labeled one hour after their terminal cell division in the Telley data, i.e., while still in the progenitor state, were most frequently expressed in the progenitors (i.e., radial glia cells) of the mouse dorsal forebrain atlas. Correspondingly, markers of 24H cells were most frequently expressed in neuroblasts (and some neurons, as noted by original authors) and markers of 96H cells in neurons of the LaManno dataset. These observations across datasets indicate that the CellCover marker gene panels from the Telley data accurately define the primary cell states traversed during neocortical neurogenesis in the mouse, as well as the significant heterogeneity in the transition between the progenitor and neuron states.

To explore if these gene panels define neocortical cell types conserved across species in mammalian neurogenesis, we also examined expression of the panels derived from the mouse in scRNA-seq data from human mid-gestational cortical tissue (Polioudakis et al. [39]; Figure 2D and 2E). Markers of 1H mouse cells are most frequently and specifically expressed in the progenitor cells of the human neocortex. Markers of 24H cells are most frequently expressed in intermediate progenitor cells (IPCs) and earliest neurons that have not migrated out of the germinal zone, while 96H markers show greatest frequency in later neurons resident in the cortical plate. This clear mapping of the murine CellCover markers onto human cortical data demonstrates that these gene panels capture neurogenic cell states conserved across mammalian development.

These transfers of the Telley 1H, 24H, and 96H CellCover marker gene panels into single-cell data of the mouse and human neocortex provide a cell-type-level perspective on the marker panels. To further understand the signals captured in these panels, we transferred them into additional data with much less cellular resolution but greater temporal and anatomical resolution. In bulk RNA-seq of human postmortem prefrontal cortical tissue [40], we observed a decrease in the expression of 1H markers across human neocortical development *in utero* (Figure 3A). This is consistent with the progressive disappearance of neural progenitors as post-mitotic neurons are generated during peak neurogenesis in mid-gestation. The expression of 24H markers appears to peak around GW12. Markers of 96H cells increase in expression during fetal development, plateauing around GW16, consistent with the progressive accumulation of newly born neurons in the developing cortex during this period.

**Figure 3:**
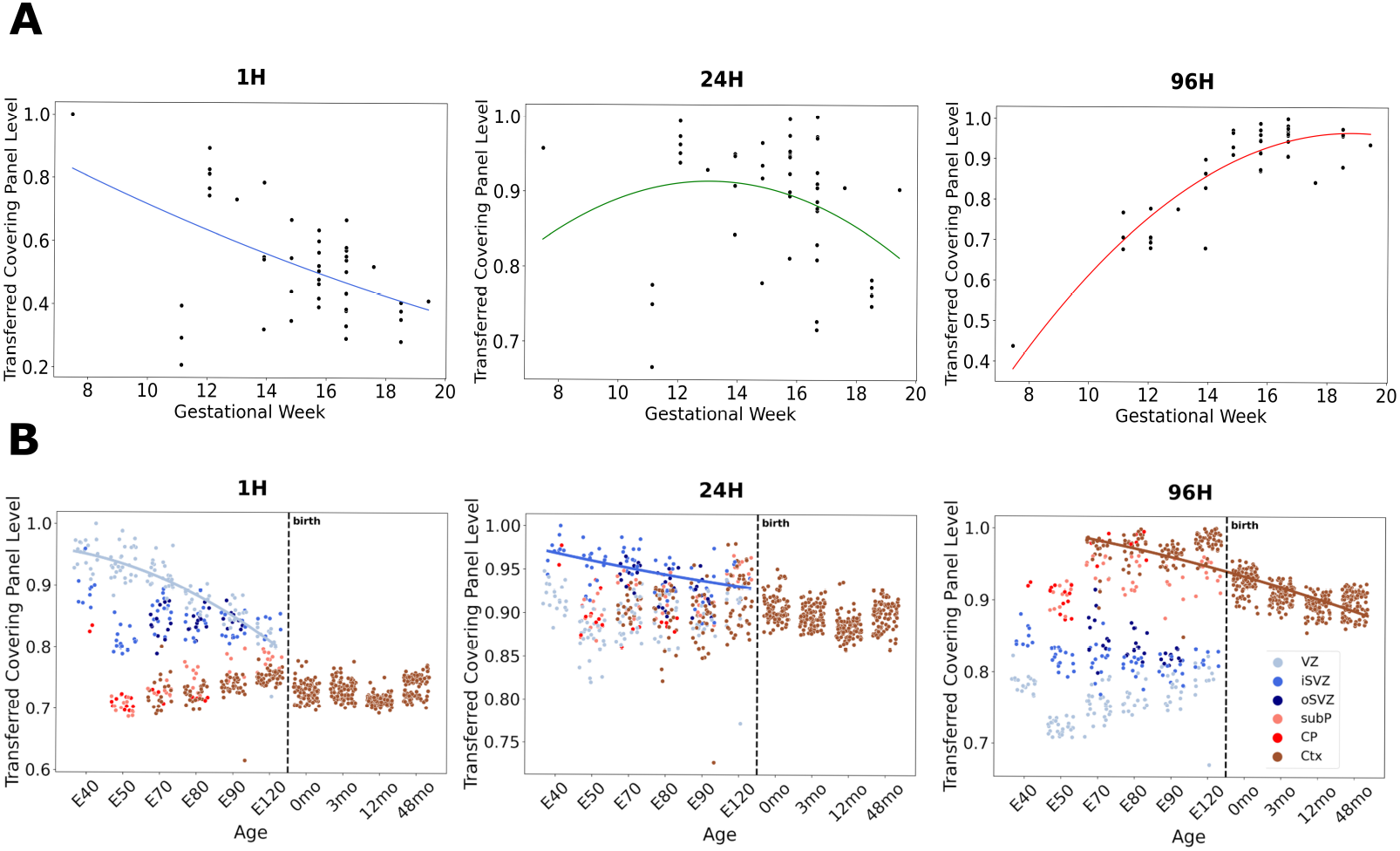
Expression of 1H, 24H, and 96H CellCover marker gene panels across development in bulk human and microdissected macaque neocortical tissue. **A**. Transfer of CellCover marker gene panels from the Telley data in mouse [34] into bulk RNAseq from human fetal cortical tissue [40]. The panel is obtained using the CellCover with ***α*** = 0.02 and ***d*** = 5. The transferred values of gene panels were assessed as the sum of panel gene expression levels in each individual sample divided by the maximum sum observed in the samples. **B**. Transfer of marker gene panels (Table S4d: ***α*** = 0.02, ***d*** = 15 nested expanded from ***d*** = 7) from the Telley data in mouse into microarray data from laser microdissected regions of the developing macaque neocortex [37]. Nonlinear fits in 1H, 24H, and 96H panels are to VZ, iSVZ, and Ctx data, respectively. X-axis ages are expressed as embryonic (E) days after conception and months (mo) after birth. Transferred marker gene panel levels were calculated as in panel A. VZ = ventricular zone, iSVZ= inner subventricular zone, oSVZ = outer subventricular zone, subP = subplate, CP = cortical plate, Ctx = cortex.

Cell states across mammalian neurogenesis are tightly linked to the position in the developing cortex. To explore the precise positional association of signals captured by CellCover panels from the developing mouse cortex, we transferred the markers derived from the Telley data into expression data from hundreds of laser microdissected regions of the developing primate cortex (Bakken et al. [37]; Figure 3B). Markers of the 1H mouse neural progenitor cells show the highest expression in the ventricular zone (VZ) of the macaque and decrease over fetal development (Figure 3B left panel), consistent with the widely held notion that ventricular radial glia are the earliest and most conserved neural progenitor state in the developing mammalian cortex. In contrast, markers of 24H cells in the mouse are most highly expressed in the cells of the inner subventricular zone (iSVZ) of the primate (Figure 3B center panel), indicating a progression from the ventricular zone state to delaminated intermediate progenitor (IPCs) / basal progenitor (BPs) states, but not reaching the primate-specific outer subventriclar zone (oSVZ) [30] / outer radial glia (oRG) states [32]. Genes marking 96H cells are most highly expressed in the postmitotic neurons of the fetal cortex and decrease following birth. Given the greater signal of the 96H gene panel in more mature neurons in Figure 2E, we conclude that these genes are not related to transient newborn neuron states. Rather, it is likely that the increasing glial content of the postnatal neocortex begins to reduce the relative level of the neuronal signal in this bulk RNA-seq data, despite its high spatial resolution. It is of interest that in the VZ, the 1H signal is high and falling while the 96H signal is low but rising. These opposing dynamic signatures within mammalian neural progenitors appear to indicate that progenitor and neuron states begin to converge over development. Elements of this have been noted elsewhere [34, 42] and become clearer later in this report as we perform additional transfers of marker gene panels at higher resolution (Figures 4 and 5).

**Figure 4:**
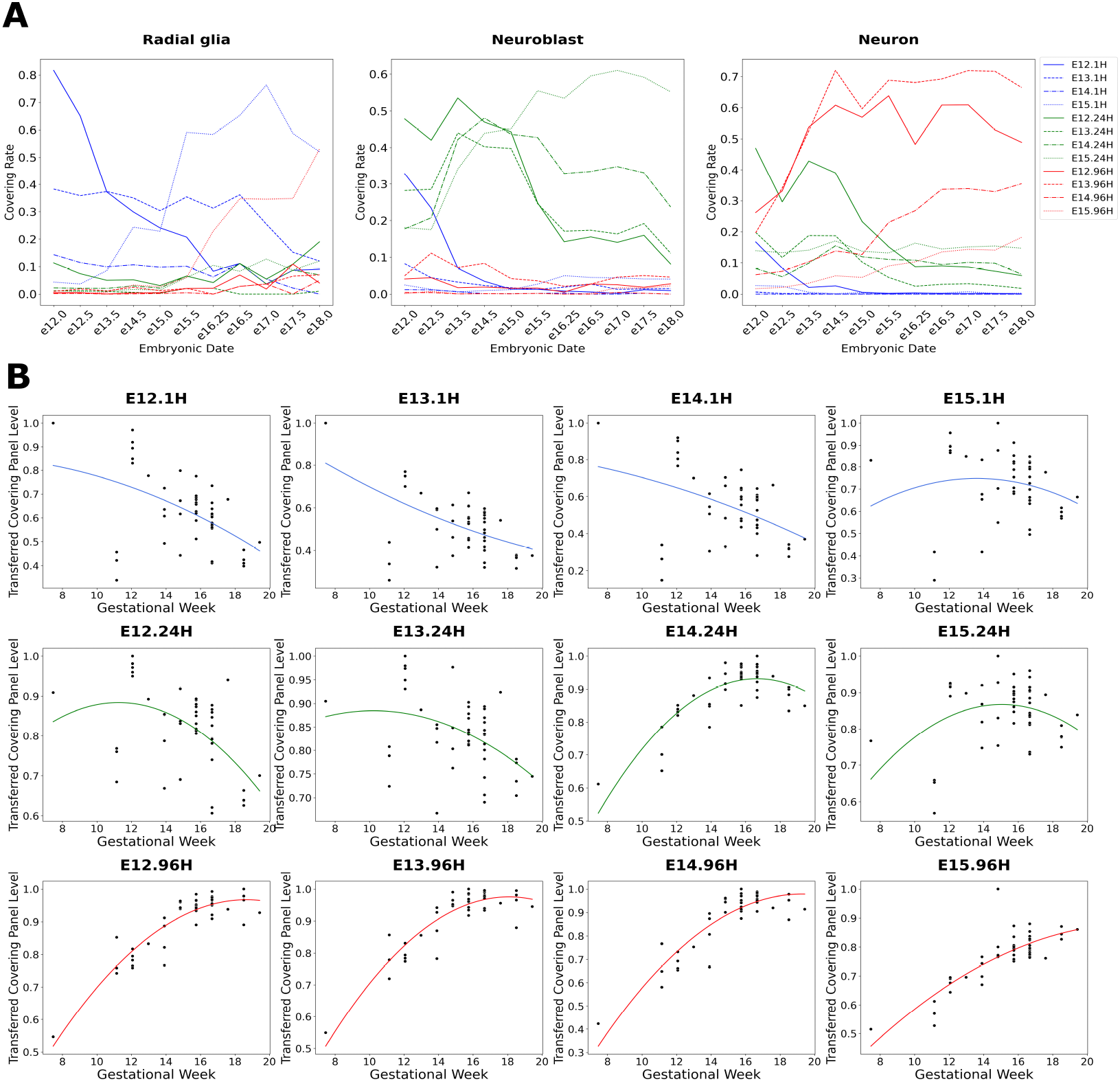
Expression of the 12 CellCover gene panels from the Telley data across development in the fetal neocortex of the mouse and human. **A**. Transfer of Telley gene panels (***α*** = 0.02 and ***d*** = 6) into the radial glia (left panel), neuroblasts (center panel) and neurons (right panel) from the developing mouse brain atlas (La Manno et al. [35]). In all cases, covering rates of transferred panels from 1H cells are depicted in blue, 24H in green, and 96H in red. Transferred levels were calculated as the proportion of cells of each type expressing more than 3 marker genes in the gene panel. **B**. Transfer of the 12 gene panels (***α*** = 0.02 and ***d*** = 15 nested expanded from ***d*** = 7) into bulk RNA-seq data from the human fetal cortex [40]. Transferred values of gene panels were assessed as the sum of gene panel expression levels in each individual sample divided by the maximum sum observed in the samples (as in Figure 3A).

**Figure 5:**
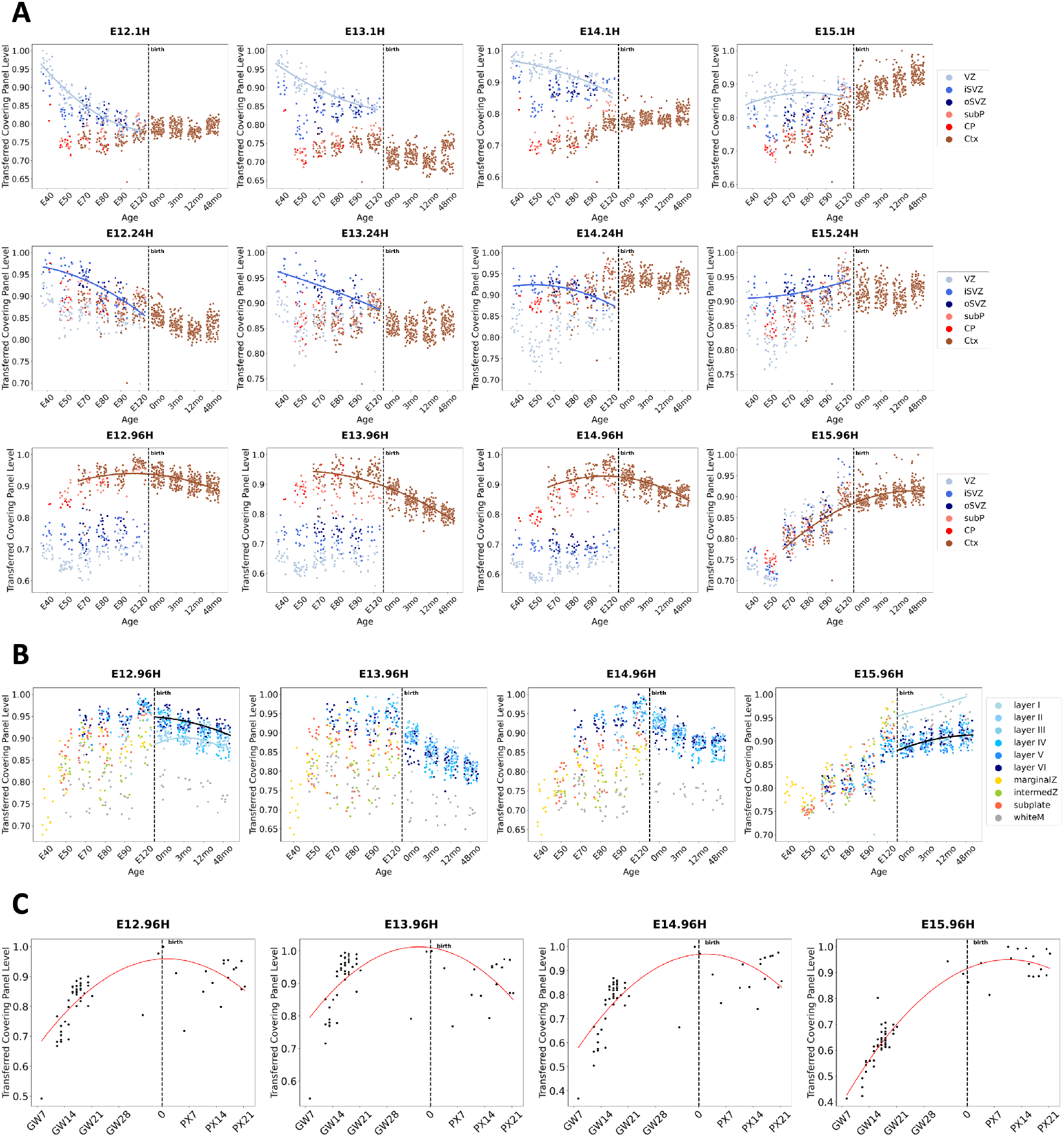
Expression of the 12 CellCover gene panels from the Telley data across development in the fetal and early postnatal neocortex of the macaque and human. The CellCover marker panels are obtained at ***α*** = 0.02 and ***d*** = 15 nested expanded from ***d*** = 7. **A**. Transfer of the 12 CellCover panels into microarray data from microdissected regions of the developing macaque neocortex [37]. X-axis ages are expressed as embryonic (E) days after conception and months (mo) after birth. Transferred gene panel levels were calculated as in Figure 3. VZ = ventricular zone, iSVZ= inner subventricular zone, oSVZ = outer subventricular zone, subP = subplate, CP = cortical plate, Ctx = cortex. **B**. Repeated transfer of the 96H gene panels into the microdissected macaque data, using additional labeling of dissections by cortical layer. **C**. Repeated transfer of the 96H gene panels into the human cortex data [40], showing additional late fetal and early postnatal samples in postnatal development.

While these concise, low covering depth gene panels can effectively map conserved cell-type-specific expression dynamics across mammalian development, the multiplicity of near-optimal solutions to the covering problem (see details in Supplemental section 3) suggests that a broader complementary approach for connecting more genes to cell classes will be useful to define underlying cell type specific mechanisms. Therefore, we proposed the CellCover nested expansion (detailed in Method section 5.2) and demonstrate its utility by expanding the 1H, 24H, and 96H gene panels to include additional marker genes. Specifically, nested expansion involves iteratively rerunning CellCover for a given cell class, using the previously optimal gene panel as a starting point and updating the algorithm with increased covering depth. We demonstrate that these expanded gene panels can be effectively used in gene over-representation analyses to explore cell function more deeply, and capture fundamentally distinct signals from scRNA-seq data than DE methods (see details in Supplemental section 4).

### 3.5 CellCover marker gene panels in mouse neocortical development #2: Cell-type-specific temporal progression

Aiming to define cell-type-specific dynamics at higher resolution, we extended our previous CellCover analysis of the Telley data from the level of three cell ages (1, 24, and 96 hours) aggregated across embryonic days (E12-15), to 12 cell class labels, each representing a single cell age from one embryonic day (3 cell ages *×* 4 embryonic days = 12 cell classes). Figure S6 depicts the individual genes in each of these 12 marker gene panels and the proportion of cells of each type which express them. Within each cell age (1H, 24H, or 96H), there is consistent expression of marker genes across embryonic days. That is, markers for cells of one age at one embryonic day are often expressed in cells of the same age at other embryonic days. This indicates common molecular dynamics in progenitors following their terminal division at the ventricle, regardless of the embryonic day on which that division took place. These shared elements are most clear in E13 cells of all ages.

In contrast, at E12, 1H cells are most distinct from other 1H cells. This particularly early progenitor state is also observable in the LaManno mouse data [35] (Figure 4A) and the Bakken macaque data [37] (Figure 5A). This E12 1H panel includes Crabp2, which marks a specific early neural progenitor state related to telencephalic regionalization in multiple primate datasets ([37] and [38]). This can be explored at: NeMO: CRABP2). At E14, 24H cells begin to become distinct from 24H cells of other embryonic ages, while 1H and 96H cells at E14 share dynamics across embryonic days. At E15, 24H and 96H cells are most distinct, paralleling the shift from deep to upper layer neuron generation (in section 3.6 we gain more granular insight into these observations as we transfer the marker gene panels into additional primate and human data). These 12 CellCover gene panels can be explored in all the public datasets we use in this report at ([NeMO: Telley 12 CellCover Panels](https://nemoanalytics.org/p?p=p&l=CellCover&c=TelleyCellCover12sets&algo=binary); these can be compared to marker panels derived from DE methods: [NeMO: Telley 12 Sets DE Genes](https://nemoanalytics.org/p?p=p&l=CellCover&c=TelleyDEG12sets&algo=binary)).

In additional analyses, we compared the 12 CellCover marker gene panels to marker genes defined by DE methods. Consistent with our findings in blood-derived data (Figure 1B), genes identified by the two methods are highly divergent. Individual genes selected by CellCover have lower sensitivity but higher specificity than DE-defined marker genes. Because CellCover can borrow power across genes by optimizing the panel rather than individual markers, CellCover panels achieve high sensitivity despite the lower sensitivity of individual genes that it selects, yielding complementary combinations of genes to define cell classes (see details in Supplemental section 5).

### 3.6 Transfer of CellCover marker gene panels #2: Conserved cell-type-specific temporal progression in neocortical development

Using CellCover marker gene panels for the three principal cell ages in the Telley data, we have shown that CellCover can precisely define the major cell classes in mammalian neurogenesis across multiple scRNA-seq data sets (Figures 2 and 3). To evaluate the ability of CellCover marker gene panels to delineate temporal change within these cell types across cortical development, we have conducted similar transfer experiments using CellCover marker panels for all 12 cell classes in the Telley data as described above (Figure S6). Figure 4A shows the transfer of these 12 CellCover panels into the LaManno 2021 mouse brain scRNA-seq atlas data [35] using covering rate in each cell type as a measure of the strength of mapping of the gene panels (see Method section 5.4). As expected for each of the three main cell types across neurogenesis: (1) radial glia in the LaManno data show the highest covering rates of gene panels derived from 1H cells in the Telley data (Figure 4A, left panel, in blue), (2) neuroblasts in LaManno have highest covering rates of 24H markers (Figure 4A, center panel, in green), (3) neurons have the highest rates of 96H markers (Figure 4A, right panel, in red).

Importantly, in the radial glia of the LaManno data, the four 1H cell class panels spanning E12-E15 display sequentially ordered temporal patterns across developmental ages (sequential peaks of blue lines in Figure 4A, left panel). Notably, radial glia beyond E15 in the laManno data begin to express neuronal (96H) markers. This is consistent with conclusions of Telley et al. [34] and and a previous report of ours [42], which note that late neural progenitors begin to express markers of postmitotic neurons. Here, we extend this observation by showing that this is not the emergence of a non-specific neuronal identity but rather that late neural progenitors begin to express markers specifically of the later-born neurons they are about to produce (Figure 4A, left panel, E15.96H markers in the red dotted line).

While 24H markers show highest covering rates in neuroblasts of the laManno data (Figure 4A, center panel, in green), neurons also show notable levels of these markers (Figure 4A, right panel, in green). This is consistent with the annotation by Telley et al. [34] of both progenitor and neuronal sub-populations within their 24H cells. That is, 24 hours following terminal division, some neural progenitors have progressed to an early neuronal state, while others have not fully exited the progenitor state (denoted as basal progenitors by Telley et al. [34]). It is also important to note in this analysis that Eday labels from the Telley data indicate the age of animals when cells were labeled. Hence E12.96H cells were labeled on E12 but harvested and sequenced on E16. This explains the later, and still clear, temporal progression of the Telley 96H marker gene panels in the neurons of the laManno data (Figure 4A, right panel, in red).

We also transferred the CellCover marker panels from the twelve Telley cell classes into bulk RNA-seq data from the developing human neocortex [40] (Figure 4B). Levels of the 1H gene panels from E12-14 are highest at the earliest time point (at GW8) and descend through the second trimester, indicating peak expression of these panels lies prior to the window of human cortical development studied here (Figure 4B, top row of panels). The gene panel from 1H cells derived from E15, in contrast, peaks at GW14, indicating that the progenitor state captured in the mouse E15.1H cells is conserved as a more advanced state in the human, potentially indicating a shift to gliogenesis. Transferred expression levels of the 24H gene panels all peak in the window of development captured in the human data, with panels from E12 and E13 peaking at GW12 before the E14 and E15 panels peak at GW16 (Figure 4B middle row of panels). These sequential dynamics demonstrate a parallel between the temporal progression of delaminating ventricular progenitors or basal progenitors (BPs) in the mouse and human which map to specific ages in both species: the shift from E12/13 to E14/E15 BPs in the mouse corresponds to a change in human BPs that occurs between GW12 and GW16. In a trend inverse of that observed in the transfer of the 1H gene panels, the 96H panels show coordinated increases from GW12-16 and appear to begin to plateau after GW16 (Figure 4B, bottom row of panels). Consistent with a later peaking state, there is less plateauing of the E15.96H gene panel signal in the human data.

Transfer of the 12 mouse CellCover panels into laser microdissected tissue from the developing macaque [37] recapitulated broad laminar trends that we observed in the transfer of the aggregated marker gene panels (1H panels with high expression in the cells of the VZ, 24H panels highest in the iSVZ, and 96H panels in the cortex; Figure 3B). This more detailed twelve-panel transfer experiment also revealed conserved temporal elements of neocortical development (Figure 5A, note the progressively later shifts in nonlinear fits for the E12–E15 marker sets). Consistent with the well-known “inside-out” (deep to upper layer) progression of neocortical neurogenesis, the E12.96H panel signal is highest in deep layers, while the E15.96H panel is elevated specifically in the most superficial and latest-born neurons of the cortex (Figure 5B, nonlinear fits).

Additionally, this E15.96H panel has a particularly distinct pattern, not neuron-specific like the earlier 96H panels, but rather a pan-cellular signature of the maturing cortex (Figure 5A, far right in bottom row of panels). Transfer of the E15.96H panel into additional late fetal and early postnatal data from the bulk human neocortical study [40] shows that, unlike the earlier E12–E14.96H panels, expression of this panel continues to increase dramatically in the third trimester and after birth (Figure 5C). These observations may reflect the shift from neurogenic-only states at E12–E14 to the onset of gliogenesis at E15. Further indicating that E15.96H gene panel has captured the beginning of gliogenesis, levels of E12, E13 and E14 96H gene panels are lowest in white matter, while the E15.96H panel shows high levels in this glia-rich tissue (Figure 5B, gray points). Similar to this 96H signal at E15, the E15.1H gene panel also shows increases in cortex postnatally that are also likely attributable to the generation of astrocytes, which maintain a portion of the neural progenitor transcriptome (Figure 5A, right-most panel in the top row and Figure S12). This indicates that both the progenitor (1H) and differentiated (96H) cell states contain transcriptomic elements that have begun to shift from neuro-to glio-genesis at E15.

These transfers of the CellCover gene marker panels map the temporal progression of cell identities we have learned from the mouse data onto neocortical development in the primate and human. This approach to defining conserved molecular progression in shared mammalian neural progenitor populations may help elucidate how neurons sharing defining molecular, morphological, and physiological features are produced contemporaneously in the mammalian cortex. We propose CellCover coupled with transfers of this kind as a general tool in mapping cell-type-specific gene expression signals across experimental systems, developmental time, and species.

### 3.7 Transfer of CellCover marker gene panels #3: Evolution of outer radial glia cells

While the late neocortical progenitor switch from neuro-to glio-genesis is conserved across mammals, there have been significant cellular innovations in neocortical development since the rodent-primate divergence ∼100 million years ago [43]. Outer, or basal, radial glia (oRG) are a neocortical stem cell type that has been greatly amplified in the primate and human lineages and has been implicated in the neocortical and cognitive expansion observed in gyrocephalic species ([32, 44, 45]). Here we examine the expression of CellCover marker gene panels derived from sorted cell types of the human fetal cortex to explore the evolution of this key cell type in cortical development, focusing on late progenitor cell types. Applying CellCover to scRNA-seq from sorted gliogenic progenitors and oRG cells of the fetal human telencephalon [46], we generated marker gene panels for these two distinct late human neocortical progenitor types. To understand their evolutionary relationship across mammals, we examined the expression of these markers across fetal development of the neocortex in human [47], macaque [38], and mouse [36] scRNA-seq data (Transfer of these gene panels into the many public datasets we use here can be explored online at: [NeMO: Sorted Brain Cell CellCover Panels](https://nemoanalytics.org/p?p=p&l=CellCover&c=LiuCellCoverSorted&algo=binary)).

As expected of markers of these late-stage progenitors, the proportion of markers expressed in progenitors rose across developmental time in macaque and human progenitors (Figure 6A & B). In both primate and human progenitors there are two clear cell populations which express these genes: one with expression of more oRG markers, and a later population with expression of more gliogenic markers (See Figure S23 & S24 for expression of these two marker panels in human and macaque progenitor cell subtypes). This is consistent with the current understanding of the sequential emergence of these distinct cell types in the primate and human brain. The increase in expression of human oRG markers over time was also observed in progenitor cells of the mouse neocortex (Figure 6C & Figure S22), albeit at lower proportions than in primate and human progenitors. This is consistent with the previous observation of oRG-like cells at low numbers in the developing murine neocortex [31]. Importantly, unlike in primates and humans where oRG markers are expressed in neural progenitors before the arrival of gliogenic progenitors (Figure 6A & B), all mouse progenitors which show expression of human oRG markers also show expression of gliogenic markers (Figure 6C & Figure S22). This is indicated by the absence of a cell population below the identity line, while there is a clear gliogenic population above the identity at later ages. This suggests that there are likely very few, if any, bona fide oRG cells in the developing mouse neocortex and progenitors expressing human oRG markers appear late in the mouse neocortical development and are likely gliogenic. This suggests an evolutionary model in which a portion of the oRG transcriptomic program came into being within gliogenic progenitors before the rodent-primate divergence, while additional transcriptomic programs and the bona fide oRG cell type emerged thereafter. This evolutionary model is consistent with the developmental conclusion made by the authors of the human neural progenitor sorting report who assert that gliogenic precursors are likely derived directly from oRG cells [46] and also consistent with our recent identification of a partially conserved oRG transcriptomic program with gliogenic elements that we defined using joint matrix decomposition across scRNA-seq data from these same three species [1].

**Figure 6:**
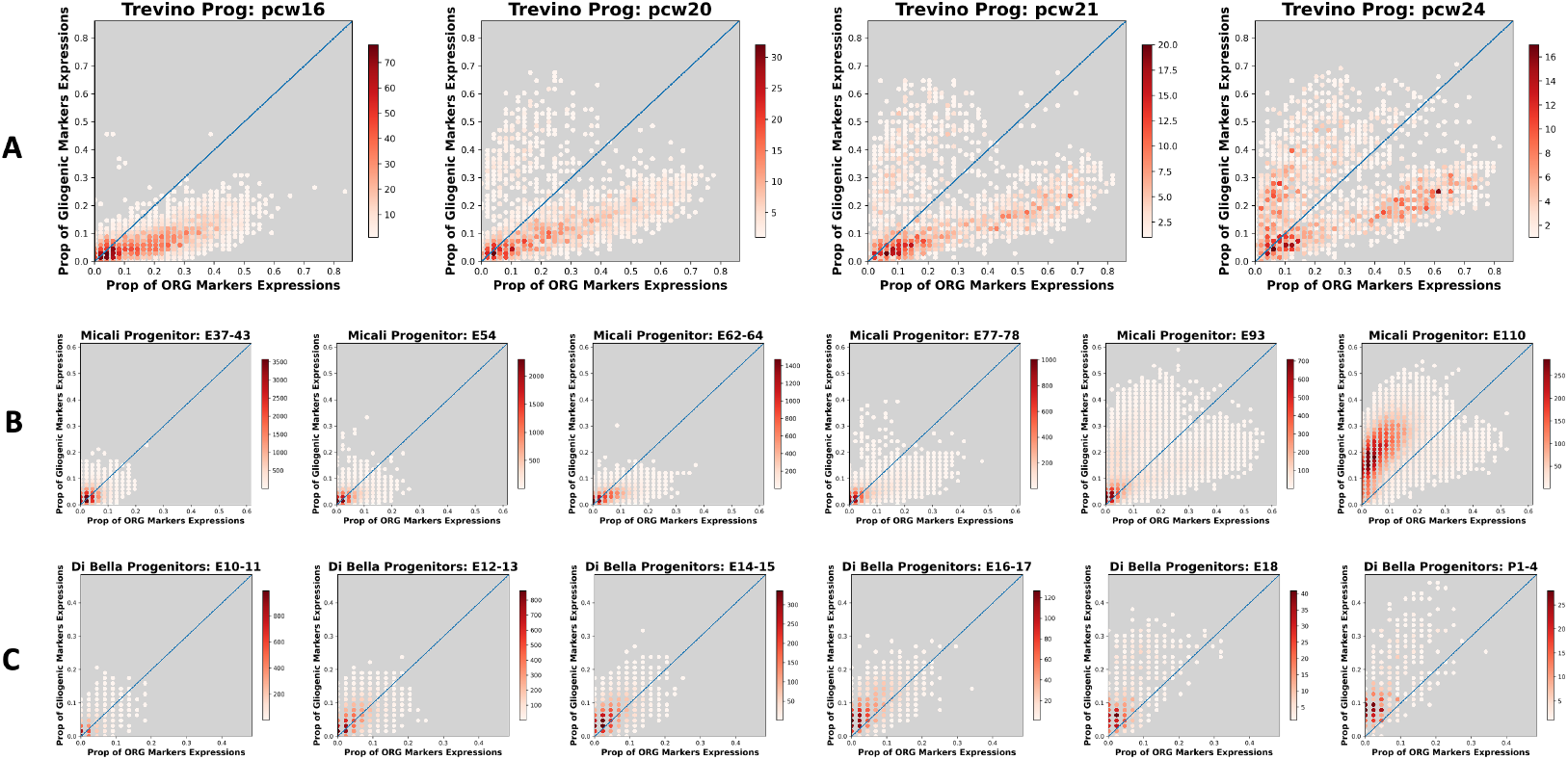
Expression of gliogenic and oRG marker gene panels across developmental time in human, macaque, and mouse neural progenitor cells. CellCover (***α*** = 0.02, ***d*** = 15) was used to define marker gene panels from scRNA-seq of sorted cell types of the developing human telencephalon [46]. Here, the expression of marker gene panels from gliogenic precursor cells and outer radial glial (oRG) cells is examined in additional scRNA-seq data from progenitor cells of the developing **A**. human [47], **B**. macaque [38], and **C**. mouse [36] neocortex. Each panel depicts the number of progenitor cells (color intensity) expressing differing proportions of oRG (X-axis) and gliogenic precrsor (Y-axis) marker panel genes at one developmental time in each species. Changes in the distribution of progenitor cells expressing different proportions of the marker genes as development progresses can be seen across panels from left to right. Ages are shown in individual panel titles. pcw = post conceptional weeks.

## 4 Discussion

In this work, we introduced CellCover, a novel algorithm designed for identifying cell-type-specific marker gene panels in scRNA-seq data. CellCover employs a minimal partial set covering approach to efficiently capture a concise set of cell type differentiating genes whose complementary expression patterns effectively distinguish cell populations. We show that marker panels selected by CellCover achieve predictive performance comparable to those identified through differential expression (DE) and other supervised marker selection methods. Importantly, CellCover consistently selects genes with less redundancy across cell types when compared to DE methods. In addition, although CellCover shares conceptual similarities with PhenotypeCover, as both adopt variants of the set covering problem, they differ substantially in objective, algorithmic formulations, and optimization procedures; a detailed comparison is provided in Supplementary section 6.

A key component of CellCover is the covering rate, which quantifies how completely a gene panel represents a cell population and, thereby, facilitates the assessment of marker gene panel generalizability across datasets. Specifically, marker panels generated from one dataset can be evaluated by their covering rate in cell populations from different datasets, enabling cross-study mapping of cell-type-specific signals. In both validation and exploration transfer experiments, we observed consistent alignment of cell types between source and target datasets along corresponding hematopoietic and neural lineages. In data from the developing mammalian neocortex, we have shown that CellCover captures cell-type-specific signals in mouse that are conserved across primate and human cortical development. Using ground truth temporal labeling of cells in the mouse cortex, we have mapped the temporal progression of murine neocortical development onto non-human primate and human developmental trajectories. Applying CellCover to sorted cell types of the fetal human cortex and transferring the resulting marker panels to mouse and primate data, we have shown that human oRG cells likely evolved through expansion of transcriptomic programs in rodent gliogenic precursors.

A limitation inherent to our method is that the partial set covering problem in the CellCover formulation is NP-hard. Consequently, integer programming solvers may experience prolonged runtimes when searching for the globally optimal solution, particularly as the required covering depth increases, e.g., when selecting a large gene panel. To address this issue, we introduced the nested expansion algorithm, in which the solution obtained at a smaller covering depth is carried forward as a subset of the solution at the subsequent larger depth. This iterative, nested approach significantly improves computational efficiency, especially when starting with a small initial covering depth and incrementally increasing it. We provide a runtime analysis of CellCover along with general implementation guidance in section 7 of the Supplementary Materials.

We invite researchers to interrogate all the public data resources we examine here in the NeMO Analytics multiomics data exploration environment that is designed to provide biologists with no programming expertise the ability to perform powerful analyses across collections of data in brain development [1]. Users can visualize individual genes ([NeMO: Individual genes in cortex](https://nemoanalytics.org/p?&l=CellCover&g=EOMES) and [NeMO: Individual genes in blood](https://nemoanalytics.org/p?&l=immune&g=CSF1R)) or groups of genes simultaneously across multiple datasets, including automated transfer of the CellCover gene marker panels we have identified across cell types and developmental time ([NeMO: Telley 3 CellCover Panels](https://nemoanalytics.org/p?p=p&l=CellCover&c=TelleyCellCover3sets&algo=binary), [NeMO: Telley 12 CellCover Panels](https://nemoanalytics.org/p?p=p&l=CellCover&c=TelleyCellCover12sets&algo=binary), [NeMO: Sorted Brain Cell CellCover Panels](https://nemoanalytics.org/p?p=p&l=CellCover&c=LiuCellCoverSorted&algo=binary), and [NeMO: Blood 34 CellCover Panels](https://nemoanalytics.org/p?p=p&l=immune&c=CellCoverHao34&algo=binary)). CellCover is available in [CellCover R](https://github.com/lanlanji/CoveringPackage) and [CellCover Python](https://pypi.org/project/CellCover/).

## 5 Methods

### 5.1 Computing marker sets

#### 5.1.1 Goals

The goal of our method is to find a relatively small set of genes for each cell type that is sufficient to label each individual cell. We associate to the observation of a (random) cell of a certain tissue a random variable *X* = (*X*_*g*_, *g* ∈ *G*) where *X*_*g*_ denotes the raw count expression level of gene *g*. We denote the cell type labels of cells by a categorical random variable *Y* ∈ {1, 2, … *K*}.

Fix a target cell type *k*. We will declare that a set of genes *M* provides a good marker set for this cell type if (1) with high probability, a sufficient number of genes in *M* are expressed if the cell type is *k*, and (2) genes in *M* are expressed with low probability if the cell type is not *k*. We note that specifying *M* is equivalent to specifying a family of binary variables *z* = (*z*_*g*_, *g* ∈ *G*), taking *M* = {*g* : *z*_*g*_ = 1}.

We will dedicate most of this discussion to the formulation of optimization problems that quantify conditions (1) and (2). We note that these conditions differ from what is most often accepted as a definition of a marker gene, a gene which is expressed with a significantly larger probability in type *k* than in any other types, or, alternatively, a gene that is differentially expressed. Such criterion defines markers at the individual gene level. However, our definition of *marker sets* relies on the collective behavior of the genes selected in *M*.

We now make more precise our definition of marker sets, which will take us to an integer linear problem (ILP).

#### 5.1.2 Binarization

We define a gene *g* to be expressed if and only if *X*_*g*_ *>* 0. This operation can be refined by replacing the zero threshold by some low activity number *θ* (or *θ*_*g*_: *X*_*g*_ *> θ*_*g*_). Given this binarization rule, we introduce the random variable *U*_*g*_ to encode the binary expression level of gene *g*, where 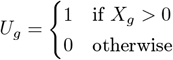.

#### 5.1.3 First condition

We now formulate condition (i) as a constraint on the set *M* and introduce two parameters of our method, namely an integer ***d*** (or “depth”) and a scalar 0 ≤ ***α*** ≪ 1 to quantify this condition as “the probability that at least *d* genes in *M* are expressed given that the cell type is *k* is larger than 1 − ***α***,” or

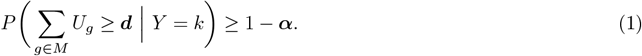

The left-hand side of the inequality (1) is the rate at which the set *M* covers the class *k*, or covering rate. Obviously, this constraint can be satisfied for some *M* ⊂ *G* if and only if it is satisfied for *M* = *G*, so that ***α*** needs to be chosen such that

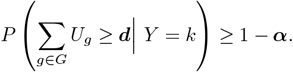

Note also that, using the notation introduced in the previous section,

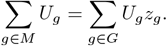

This condition can also be interpreted in terms of a classification error. Indeed, define Φ_*M*,***d***_ as the binary classifier equal to one if 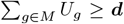 and to 0 otherwise. Then (1) expresses the fact that this classifier’s sensitivity is at least 1 − ***α***.

#### 5.1.4 Second condition

Having expressed the first condition as a constraint, we now express the second one as a quantity *M* ↦ *F* (*M*) to optimize. Ideally, one would like to minimize the probability that, conditionally to *Y*_*k*_ = 0, there exists at least one active gene in *M*, namely

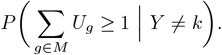

This quantity, however, is not straightforward to minimize, but its Bonferroni upper bound

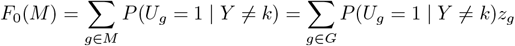

is linear in the binary vector *z* and will be amenable to an ILP formulation. (*F*_0_(*M*) is also equal to the expectation of 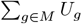 given *Y* ≠ *k*.)

Note that, obviously

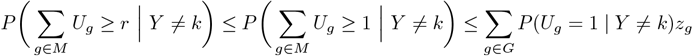

so that 1 − *F*_0_(*M*) controls the sensibility of the previous classifier Φ_*M,r*_.

If the variables *U*_*g*_ are independent (conditionally to *Y*), then

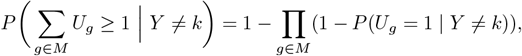

which suggests using

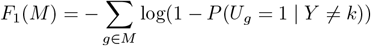

as an alternative objective.

#### 5.1.5 Margin weights

More generally, we will consider objective functions *F* taking the form

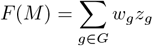

for some choice of weights (*w*_*g*_, *g* ∈ *G*). The function *F*_0_ above uses *w*_*g*_ = *P* (*U*_*g*_ = 1 | *Y* ≠ *k*), and *F*_1_ uses *w*_*g*_ = − log(1 − *P* (*U*_*g*_ = 1 | *Y* ≠ *k*)), but other choices are possible. We will in particular work with weights reflecting the ability of the variable *U*_*g*_ to identify class *k*, such as

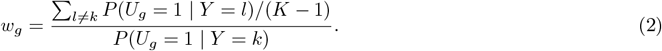

Or

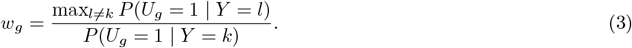

where *K* is the total number of classes. Small values of *w*_*g*_ reflect a large sensibility of the variable *U*_*g*_, when differentiating type *k* from any other type. In our algorithm, we implement Equation (2) as the default while offering users the flexibility to choose their desired weight schemes. Going further, it is natural to eliminate genes for which one of the ratios above is larger than 1, which is achieved by enforcing *z*_*g*_ = 0 if *w*_*g*_ *>* 1 (or, equivalently, replacing *w*_*g*_ by +∞ when *w*_*g*_ *>* 1). This choice has, in addition, the merit of reducing the number of free variables *z*_*g*_ and accelerating the optimization process. Note that these margin weights ensure that *F* (*M*) ≥ *F*_0_(*M*) and therefore also control sensibility.

#### 5.1.6 Marker set estimation algorithm

We can now conclude our discussion with a description of our algorithm. Assume we observe the raw count single-cell gene expression data of *N* cells 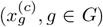 and their cell types *y*^(*c*)^ for *c* = 1, …, *N*. We binarize 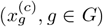 to obtain the binary expression levels 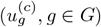 for *c* = 1, …, *N* as the input for our algorithm. Let *S*_*k*_ be the set of cells with type *k* for *k* = 1, 2, …, *K*. Given the hyperparameters **d** and ***α***, the following algorithm returns a covering marker panel for cell type *k*, with covering depth ***d*** and covering rate 1 − ***α***:

1. Estimate probability that gene *g* is expressed under cell type *l* with

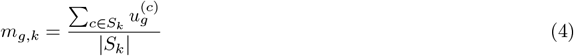

for *g* ∈ *G* and *k* = 1, …, *K* based on training data.
2. Compute weights *w*_*g*_, *k, g* ∈ *G* as the ratio between the average probability of expression of *g* when cell type is not *k* and the probability of gene *g* expressed under cell type *k* (or use user-supplied weights).

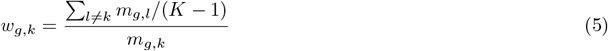 A small value of *w*_*g,k*_ indicates that gene *g* is discriminative of cell type *k*.
3. To form the candidate gene set *G*_*k*_, filter out the genes that are either not discriminative or too lowly expressed, letting *G*_*k*_ = {*g* : *w*_*g,k*_ ≤ 1} ∩ {*g* : *m*_*g,k*_ ≥ 0.1}.
4. Use an integer programming software [48] to minimize, with respect to binary variables *z*_*g*_, *g* ∈ *G*_*k*_ and *s*^(*c*)^, *c* ∈ *S*_*k*_, the function 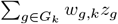 subject to the constraints

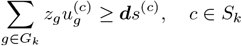

and

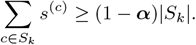

where
  - *z*_*g*_ is the marker gene indicator variable such that *z*_*g*_ = 1 if gene *g* is selected to the marker panel *M*_*k*_.
  - *s*^(*c*)^ is the cell covered indicator variable such that *s*^(*c*)^ = 1 if there are greater than or equal to **d** genes in *M* being expressed in cell *c*.
5. Return the covering panel *M*_*k*_ = {*g* ∈ *G*_*k*_ : *z*_*g*_ = 1}.

### 5.2 Nested marker set expansion

Step (4) can be modified to estimate a covering as a superset of a previous one (obtained e.g., with a smaller depth): Assume that 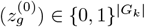 are given, e.g., from a previous run of our algorithm, defining an initial marker panel. In order to extend this panel to a covering at depth ***d***, one only needs to replace step (4) by the following: (4’) Use an integer programming software to minimize, with respect to binary variables *z*_*g*_, *g* ∈ *G*_*k*_ and *s*^(*c*)^, *c* ∈ *S*_*k*_, the function 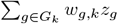 subject to the constraints

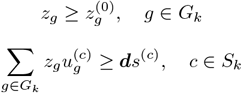

and

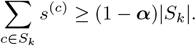

#### 5.2.1 Remarks

(a)Even though it was not designed with this intent, our marker set estimation algorithm can be interpreted as an asymmetric classification method to separate type *k* from all other types. In an approach akin to classical statistical testing, we determine a classifier with some free parameters (here, the gene set *M*) that we require to have a minimal sensitivity, 1 − ***α*** (so that ***α*** is reminiscent of a type-I error). The free parameters are then optimized in order to minimize an objective function that controls (among other things) the sensibility of the classifier.

(b)If we take *w*_*g*_ = 1 for all *g*, the previous algorithm only depends on training data associated with cell type *k*. The resulting algorithm provides a *minimal* gene set that covers the considered population, in the sense that all cells in that population (except a fraction ***α***) has at least ***d*** active genes in the selected set. With a suitable definition of what is meant by being active, this covering algorithm was introduced in Ke et al. [19] and applied to the determination of important gene motifs in a population of tumor cells associated with a specific phenotype.

### 5.3 Benchmark Analysis

#### 5.3.1 Support Vector Machine

To evaluate the performance of marker genes identified by different methods, we employed a Support Vector Machine (SVM) classifier using the *sklearn*.*svm*.*SVC()* function from the scikit-learn library [49]. SVMs are supervised learning models that construct optimal hyperplanes in high-dimensional spaces to effectively separate multiple classes with maximum margin. To address class imbalances in our multi-class gene expression data, we applied class weight normalization by setting the *class*_*weight* parameter to “*balanced*^*′′*^. This approach assigns weights to each class inversely proportional to their frequency, ensuring equal representations of different cell types during the training process. The SVM was trained exclusively on gene expression data restricted to the selected marker genes, allowing us to assess the discriminative power of these markers in accurately predicting class labels across different methods.

#### 5.3.2 Balanced Accuracy

Balanced Accuracy was employed as the primary performance metric to mitigate the effects of class imbalance in our multi-class classification. Unlike conventional accuracy, which may overemphasize performance in the more prevalent classes, balanced accuracy measures the average recall (also referred to as sensitivity) across all classes. Specifically, for each class, the true positive rate is calculated, and these rates are then averaged over all classes.

#### 5.3.3 Gene Weight Computation Based on Binary and Log-normalized Expression Profile

In the benchmark analysis, CellCover employs two weighting schemes for gene selection. The first uses the default weight defined on the binarized gene expression data (Equation 4 in Methods Section 5.1.6), while the second modifies this approach by using log-normalized gene counts in place of binary expressions:

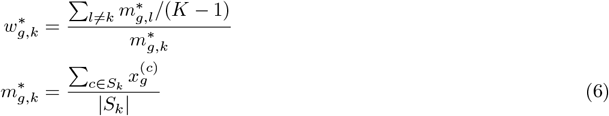

where 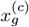 is the log-normalized expression of gene *g* in cell *c*. In both cases, the covering constraints are consistently defined using the binarized data. These two weighting schemes were implemented to ensure fair comparisons with other methods, as RankCorr, scGeneFit, PhenotypeCover, and DE all use log-normalized count data as input by default. Throughout the paper, unless otherwise specified, we use binary data for both weight calculation and covering constraints in CellCover.

#### 5.3.4 Intersection of Global Marker Panels

We compare each method’s global marker panel to the global marker panel produced by CellCover (log-normalized). Specifically, for a given method, we count the number of genes that overlap with the CellCover (log-normalized) global marker panel and then divide by the total number of genes in the CellCover (log-normalized) global panel. This ratio yields the proportion of intersection between each method’s global marker panel and the CellCover (log-normalized) reference panel.

#### 5.3.5 Within Marker Panel Redundancy

We further assess the redundancy of marker genes within the global marker panel identified by each method *x* ∈ {CellCover-LogNorm, DE}. For each cell type *k* ∈ 1, …, *K*, we obtain a local marker panel 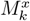 of same size. We then define the global marker panel *M*^*x*^ as the union of all cell-type-specific marker panels

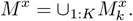

Next, let *R*^*x*^ ∈ *M*^*x*^ be the set of genes appearing in at least two different local marker panels. The proportion of redundant genes in *M*^*x*^ is thus

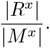

This measure reflects how frequently the same gene is identified as a marker for multiple cell types by a given method.

### 5.4 Cell Type Mapping Across Datasets

To quantify the consistency of mapping between cell types in the source and target datasets using the CellCover marker gene panels, we calculated the proportion of cells in each target cell type expressing at least 5 markers in the CellCover marker panel for each source cell type. We then applied 0-1 normalization to the covering rates of *M*_*k*_ across all available cell types *k*^*′*^ in the target dataset. This normalization was performed to standardize the covering rates of marker panel *M*_*k*_, ensuring uniform comparability across all target cell types for each source cell type marker panel. To establish a mapping between cell types in the source and target datasets, we required the normalized covering rate to exceed a threshold of 0.8.

## 6 Acknowledgements

Data sharing and visualization via NeMO Analytics was supported by grants R24MH114815 and R01DC019370. The work of DG, LY, and LJ was partially supported by NIH National Cancer Institute Grant R01CA200859. The work of DG, LY, and AW was partially supported by NSF Award 2124230.

