## Supplemental Materials for "CellCover Defines Marker Gene Panels Capturing Developmental Progression in Neocortical Neural Stem Cell Identity"

#### 1 Binarizing scRNA-seq count data

Binarization of gene expression data, where zero counts remain zero and positive counts map to 1, is a key step in defining minimal coverings of cell types with CellCover. This approach has been used in various contexts, including the study of cell-type-specific transcriptional dynamics [1], imputing the count expression profile [2], and cell age prediction [3]. We evaluated the effects of binarization on data exploration using the Telley dataset [4], which contains scRNA-seq data labeled by both the developmental age of the mouse pup (E12–E15) as well as the exact time since the terminal division of each individual progenitor cells (1, 24, or 96 hours). Throughout this report we leverage this uniquely annotated dataset to explore the developmental progression of mammalian neocortical neurogenesis. These 12 cell classes (4 pup ages x 3 times since final cell division) were analyzed to determine whether binarized data could preserve class-specific expression patterns. Two methods were applied:

(1) **Supervised prediction of ground truth labels:** Among various combinations of data types and prediction models, PCA-processed binarized data paired with a one-layer neural network achieved the highest average prediction accuracy for the 12 classes (Table S1). Similar results were observed in the Polioudakis dataset [5], where binarized data outperformed log-transformed counts in predicting cell locations (Table S2).

(2) **Unsupervised dimension reduction and clustering:** Using t-SNE or UMAP embeddings and Louvain clustering in the Telley dataset, binarized data achieved clearer separation of developmental cell classes compared to continuous counts (Figure S1). Quantitatively, binarized data produced higher adjusted Rand indices (ARIs), demonstrating stronger concordance between ground truth labels and clustering results (Table S3).

These results show that binarized gene expression data maintain sufficient structure to analyze class-specific patterns effectively.

| **XGB** | **$XGB_b$** | **$XGB^{PCA}$** | **$XGB_b^{PCA}$** | **NN1** | **$NN1_b$** | **NN2** | **$NN2_b$** | **NN3** | **$NN3_b$** |
|---|---|---|---|---|---|---|---|---|---|
| 95.0 | 93.6 | 86.5 | 91.2 | 93.8 | **97.1** | 93.3 | 96.4 | 87.9 | 94.9 |

**Supplementary Table 1**: **Average accuracies of different data types and supervised classifiers in recovering the 12 time-based cell labels in the Telley data.** Subscript "b" indicates binary data; all other trials used normalized log2 count level data. XGB = XGBOOST [6]. "PCA" indicates using only the 1st 100 principal components. NN$x$ refers to a neural network with $x$ hidden layers and 64 total nodes in the network. The size of the training set is 2342, and the size of the test set is 414.

| Donor | **XGB** | **$XGB_b$** | **$XGB^{PCA}$** | **$XGB_b^{PCA}$** | **NN1** | **$NN1_b$** | **NN2** | **$NN2_b$** | **NN3** | **$NN3_b$** |
|---|---|---|---|---|---|---|---|---|---|---|
| **368** | 91.7 | 90.4 | 91.1 | 92.7 | 91.0 | **93.1** | 90.7 | 92.6 | 90.6 | 92.5 |
| **370** | 87.2 | 84.7 | 86.7 | **87.8** | 85.9 | 87.1 | 86.6 | 87.5 | 86.2 | 87.3 |
| **371** | 91.8 | 90.8 | 90.9 | 91.8 | 89.7 | **91.9** | 90.0 | 91.8 | 90.1 | 91.6 |
| **372** | 92.0 | 91.1 | 91.7 | 92.4 | 92.1 | **92.8** | 91.9 | 92.6 | 91.7 | 92.4 |

**Supplementary Table 2**: **Comparison of binary and count data in predicative capacity in Poliousdakis dataset.** Subscript "b" indicates binary data, other trials used normalized log count level data. XGB = XGBOOST. "PCA" indicates where classification was based on the 1st 100 principal components. $NN_x$ = Neural net with $x$ layers, where $x$ indicates the number of hidden layers with 64 nodes in the network.

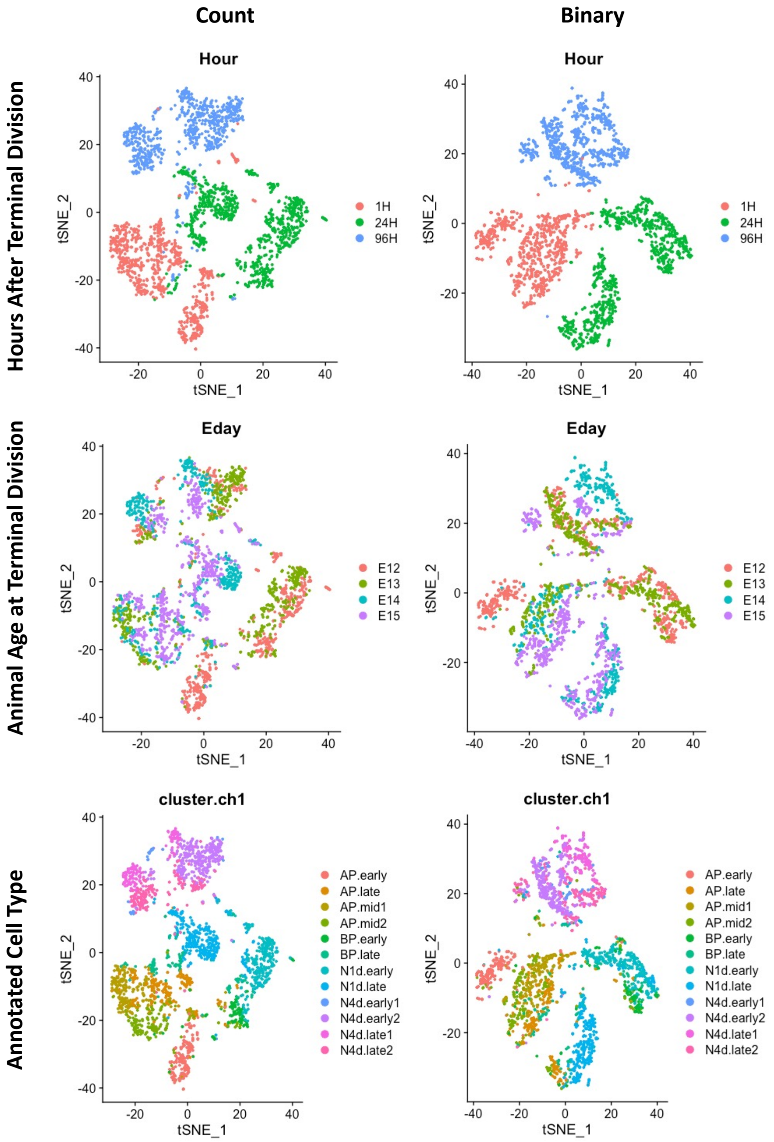


**Supplementary Figure 1**: Comparison of raw count and binarized scRNA-seq data in standard unsupervised dimension reduction using the Telley [4] dataset where ventricular neural progenitor cells were labeled specifically during their final cell division and collected cells for sequencing. To focus on precise cell types, only cells of the excitatory glutamatergic neuronal lineage were included in this analysis (this is the case in all analyses in this report).

| | **Count** | **Binary** |
|---|---|---|
| **random 2000 genes** | 0.32 | 0.37 |
| **random 4000 genes** | 0.34 | 0.39 |
| **random 6000 genes** | 0.36 | 0.41 |
| **random 8000 genes** | 0.35 | 0.44 |
| **all genes** | 0.37 | 0.45 |

**Supplementary Table 3**: **Adjusted Rand Indices (ARI) for different inputs.** Each entry of the table shows the ARI between Seurat clustering results and true time clusters for each version of input data (count or binary expression profile of the indicated number of genes).

#### 2 Benchmark

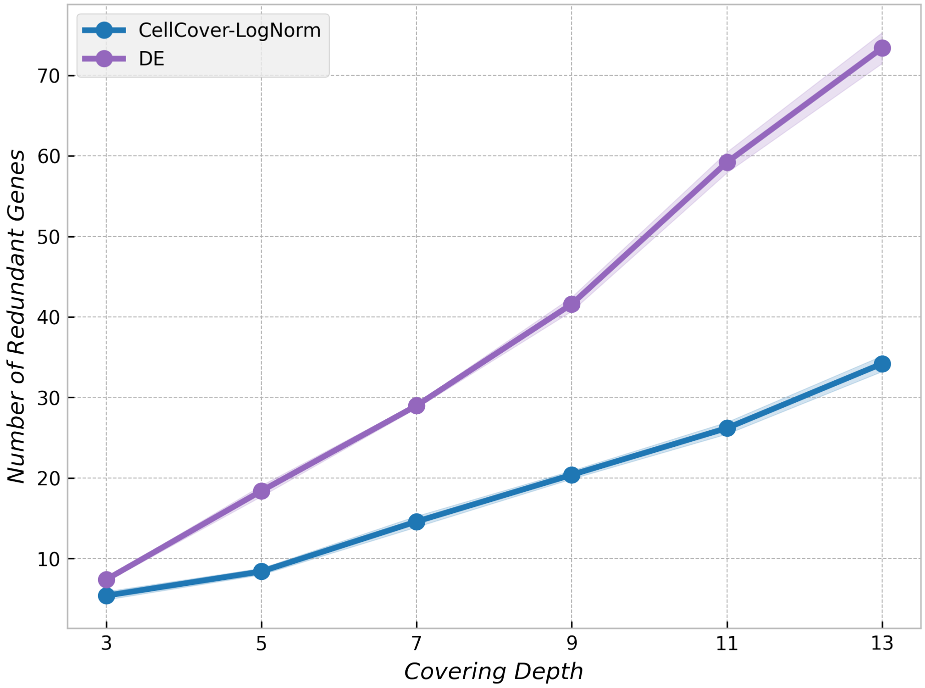


**Supplementary Figure 2**: **Number of redundant marker genes versus panel size for CellCover and DE:** The figure includes standard deviation across random seeds.

### 3 Bootstrap Analysis

The compact marker panels generated by CellCover capture cell-type specific signals in mouse neurogenesis within a dataset (Main Figure 2A), which are consistent across additional mouse data (Main Figure 2B) and are conserved across species when transferred to primate (Main Figure 3B) and human prenatal scRNA-seq data (Main Figure 2D and 3B). While these small sets of marker genes are useful for identifying cell classes, theoretically, there exist many unique minimal covering panels which satisfy a specific covering depth, $\boldsymbol{d}$ and false negative covering rate, $\boldsymbol{\alpha}$. To demonstrate this empirically, we used a nonparametric bootstrap approach whereby for each of the three cell ages in the Telley data, we sampled with replacement from the cells for the age of interest using a sample size equal to the number of cells in the original sample and again ran the CellCover algorithm (a conventional bootstrap design). This was repeated in each of 100 iterations with parameters identical to those used to define the original covering panels in Main Figure 2A. Supplementary Figure 3 lists the 15 genes which appeared in the most covering solutions along with the fraction of solutions in which they appeared. Across the covering marker gene panels for the three cell ages, only between 3 and 7 of the top 15 genes appeared in all solutions across the 100 bootstraps iterations for the 3 cell classes, and several top genes appeared in $<$50% of solutions. Similar results were observed in a bootstrap analysis of CellCover in the Geschwind data distinguishing cells captured from the germinal zones versus cortical plate (Supplementary Figure 4).

This multiplicity of good solutions to the covering problem in cell classification (and the cell type marker problem in general) indicates that while the small covering marker gene panels can effectively identify cell classes, a broader complementary approach for connecting more genes to cell classes will be useful in order to define underlying cell type specific mechanisms (see next Supplemental Section 4).

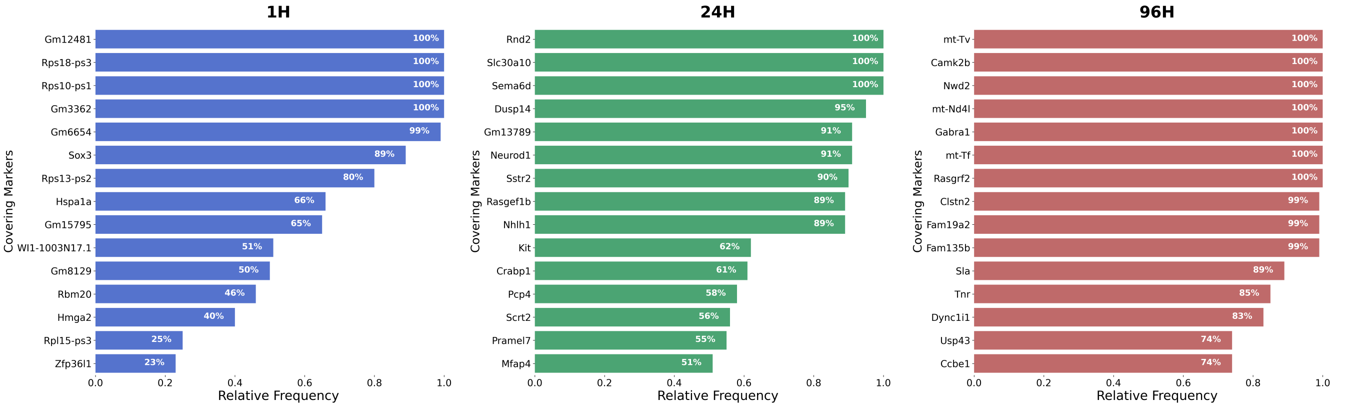


**Supplementary Figure 3**: **CellCover markers genes with highest relative frequency.** This bar plot shows the top cell age covering markers ordered by the relative frequency of each gene being selected by covering algorithm with $\boldsymbol{d} = 5$, $\boldsymbol{\alpha} = 0.02$ on the Telley dataset.

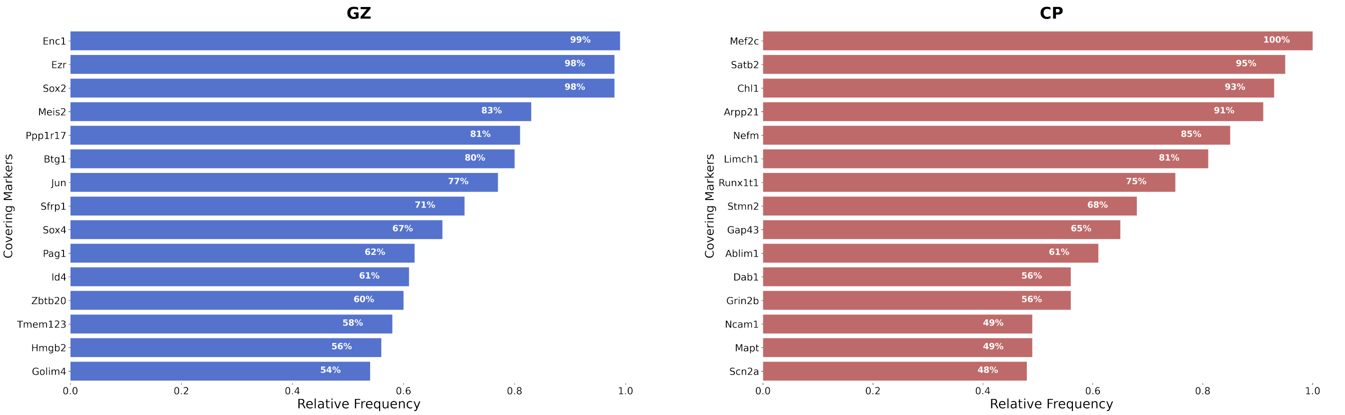


**Supplementary Figure 4**: **CellCover markers genes with highest relative frequency.** The bar plot shows the top cell age covering markers ordered by three relative frequency of each gene being selected by covering algorithm with $\boldsymbol{d} = 3$, $\boldsymbol{\alpha} = 0.05$ on the Geschwind dataset.

### 4 Using deeper covering analysis to more broadly explore cell function

We propose an additional approach in Methods section 5.2 to leverage the cell type specific signals captured by CellCover in order to extend the exploration of what cells are (classes) to include what cells are doing (cell type specific functions). From a small covering panel, this will create a broader list of genes that can be used to explore cell-type specific function via gene set over-representation analysis and more complex transfer learning experiments. To expand a covering gene marker panel for a particular cell class, the CellCover algorithm is simply run again, taking the initial covering set as a starting point along with new, broader covering constraints (depth, $\boldsymbol{d}$ and proportion of uncovered cells, $\alpha$) as input. Enforcing the inclusion of the previously defined covering panel in this nested run ensures the original compact list of genes for cell identity will be a subset of the expanded panel to be used in the exploration of cell function and also reduces the CellCover run time (compared to a de novo run with the same expanded constraints - see Methods Section 5.2).

Table S4 lists expanded covering marker gene panels for each of the three cell ages in the Telley data that we have been exploring. The original covering gene marker panels of depth 3 were expanded to depths of 7, 15, 20, 25, 30, 35, and 40. To assess the capacity of the covering marker gene panels of differing size to characterize cell type specific function, we performed over-representation analysis using Gene Ontology (GO) on all these covering panels in addition to marker panels of equal size derived from conventional DE marker selection (Figure S5). In all cases, increasing panel sizes resulted in the detection of increasing numbers of significantly enriched gene sets. The one exception to this is in the 24H panels where, after an initial increase, a reduction in the number of hits is observed in both methods, before hits again begin to rise with increasing panel size. Strikingly, while the number of significant hits for both methods grows with increasing panel sizes, the specific gene sets found to be enriched by the two methods are quite distinct: 40–50% of the gene sets detected by one method are not found to be significant by the other. This is consistent with our other observations indicating that the two methods of gene marker definition extract fundamentally different signals from single-cell data (see Supplement Section 5).

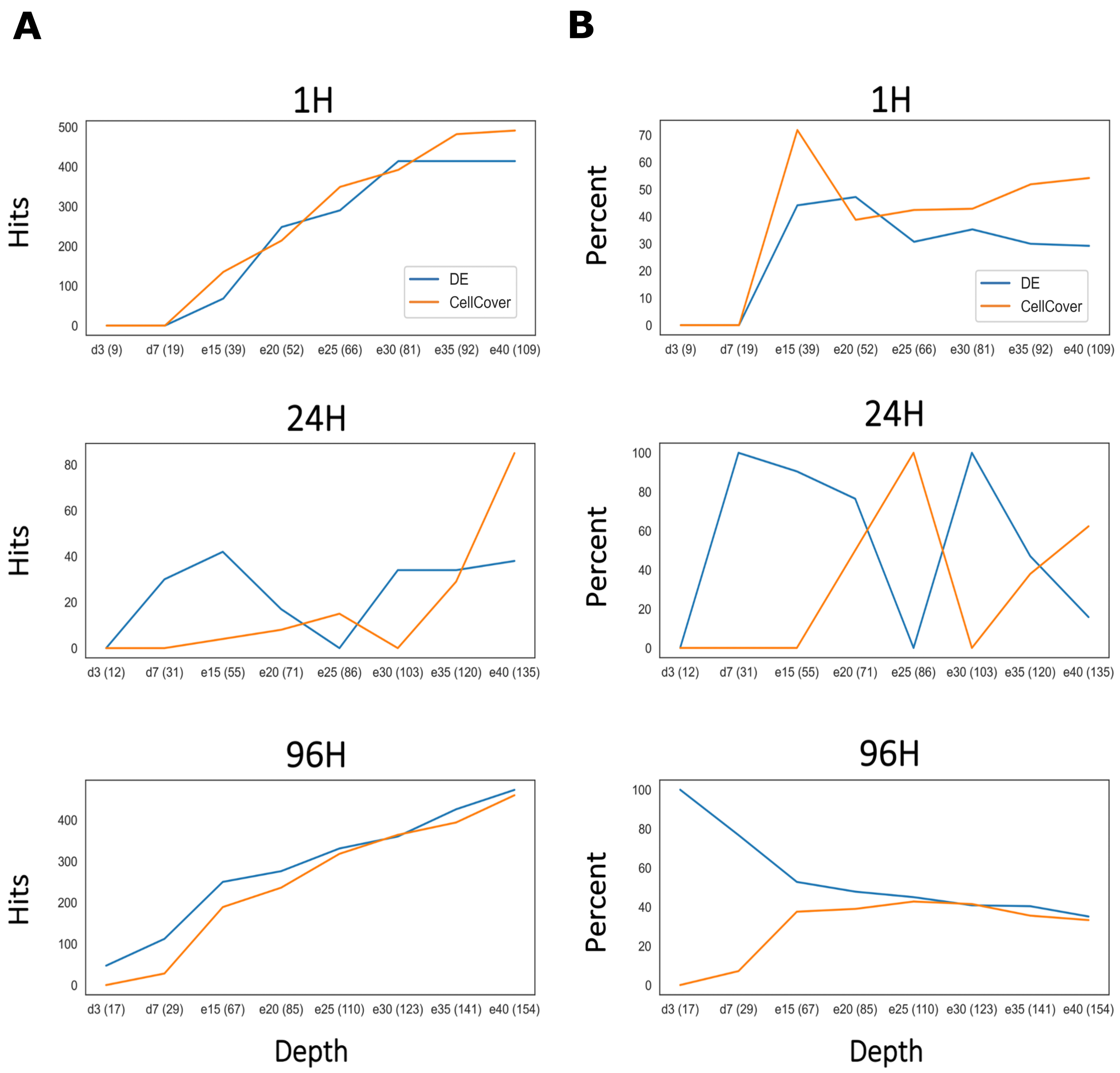


**Supplementary Figure 5**: **Over-representation analysis in gene marker panels of differing size and methodology, using the GO database.** Over-representation analysis was performed using the Fisher's exact test using gene sets from the GO database. Over-represented gene sets were marked as significant by a $FDR < 0.1$ cutoff using the Benjamini-Hochberg method. The background gene list was defined as all the genes present in the Telley dataset. **A.** The number of GO gene sets found to be significantly enriched in the gene marker panels derived from CellCover and DE methods across different matched panel sizes. The X-axis indicates the depth parameter d that was set for the CellCover run (preceding the numbers on the X-axis, "d" indicates the depth in de novo CellCover runs, "e" indicates expanded runs that use marker panels from the preceding CellCover run as a starting point). Corresponding numbers of DE markers were selected for each depth so that marker panels of equal size were compared across CellCover and DE methods. Numbers in parentheses indicate the number of genes in each marker panel. **B.** The proportion of gene sets that were found to be significant by one method and not the other.

| | |
|---|---|
| 1H | Rbm20, Gm12481, Sox3, Tcf7l1, Rps18-ps3, Gm8129, Rps10-ps1, Rps13-ps2, Gm3362 |
| 24H | Rnd2, Kit, Dusp14, Pramel7, Slc30a10, Sema6d, Sh3bgrl2, Neurod1, Nhlh1, Gm15232, Gm13789, Gm5619 |
| 96H | Gabra1, Rasgrf2, Sla, Ntsr1, Clstn2, N28178, Fam135b, Fam19a2, Ccbe1, Adap1, Camk2b, mt-Tf, mt-Tv, mt-Nd4l, Gm15344, Nwd2, Gm26871 |

(a) **Covering markers of Telley dataset with $d = 3$ and $\alpha = 0.02$**

| | |
|---|---|
| 1H | Ajuba, Etl4, Rbm20, Gm12481, Sox3, Gas1, Rps18-ps3, Gm8129, Rps10-ps1, Rpl15-ps3, Rps13-ps2, Gm6654, Gm3362, Hspa1a |
| 24H | Rnd2, Kit, Dusp14, Serping1, Ifitm3, Pramel7, Slc30a10, Sema6d, Rasgef1b, Tmem176b, Crabp1, Neurod1, Mfap4, Sstr2, Nhlh1, 9630028B13Rik, Scrt2, mt-Ti, Adamtsl3, Gm12005, Gm15232, Gm13789, Gm5619 |
| 96H | Mef2c, Gabra1, Tnr, Usp43, Rasgrf2, Sla, Dync1i1, Ptpro, Clstn2, N28178, Fam135b, Kcnj2, Fam19a2, Ccbe1, Adap1, Camk2b, Unc5c, mt-Tf, mt-Tv, mt-Nd4l, Gm15344, Nwd2, Gm26871, RP23-14P23.9 |

(b) **Covering markers of Telley dataset with $d = 5$ and $\alpha = 0.02$**

| | |
|---|---|
| 1H | Ajuba, Notch2, Rbm20, Gm12481, Sox3, Cenpe, Gas1, Tcf7l1, Hmga2, Rps18-ps3, Gm8129, Rps10-ps1, Rpl15-ps3, Sox21, Rps13-ps2, Gm6654, Gm3362, Hspa1a, WI1-1003N17.1 |
| 24H | Rnd2, Kit, Dusp14, Gadd45g, Wnt7b, Serping1, Ifitm3, Pramel7, Tuba4a, Slc30a10, Sema6d, Rasgef1b, Cabp1, Tmem176b, Sh3bgrl2, Crabp1, Neurod1, Inf2, Ttc39b, Sstr2, Plcb1, Nhlh1, 9630028B13Rik, Scrt2, Adamtsl3, Gm12005, Rpl38-ps1, Gm15232, Gm13789, Pcp4, Gm5619 |
| 96H | Mef2c, Gabra1, Tnr, Usp43, Rasgrf2, Sla, Dync1i1, Grip2, Ptpro, Clstn2, Arpp21, N28178, Fam135b, Dnm3, Rims1, Fam19a2, Ccbe1, Lingo1, Adap1, Camk2b, Unc5c, mt-Tf, mt-Tv, mt-Nd4l, Gm15344, Nwd2, Uba52, Gm26871, RP23-14P23.8 |

(c) **Covering markers of Telley dataset with $d = 7$ and $\alpha = 0.02$**

| | |
|---|---|
| 1H | Nr2e1, Zfp36l1, Spata13, Ajuba, Col2a1, Hes1, Flnb, Notch2, Cenpa, Gsta4, Aspm, Etl4, Rbm20, Gm12481, Sox3, Cenpe, Gas1, Cdca7, Dach1, Tcf7l1, Hmga2, Rps18-ps3, Gm8129, Rps26-ps1, Rps10-ps1, Rpl15-ps3, Sox21, Rpl34-ps1, Rps13-ps2, Sox2, Gm10224, Gm6654, Gm3362, Gm13267, Gm15795, Hspa1b, Hspa1a, Gm28625, WI1-1003N17.1 |
| 24H | Rnd2, Kit, Dusp14, Nuak1, Gadd45g, Lrrc16b, Wnt7b, 5330426P16Rik, Serping1, Ifitm3, Pramel7, Tuba4a, Slc30a10, Sema6d, Necab3, Rasgef1b, Cabp1, Tmem176b, Sh3bgrl2, Crabp1, Eomes, Shf, Neurod1, Osbpl5, Inf2, Ttc39b, Clvs1, Sox5, Shb, Sstr2, Bcl11b, Nhlh2, Plcb1, Nhlh1, 9630028B13Rik, Rpl31-ps13, Cntn2, A930024E05Rik, Itpk1, Scrt2, Cxcl12, Unc5d, mt-Ti, Adamtsl3, Gm23935, Gm4294, Gm12005, Gm15481, Rpl38-ps1, Gm11989, Gm15232, Gm13789, Pcp4, Rps19-ps12, Gm5619 |
| 96H | Ndrg1, Mef2c, Gabrb2, Crlf1, Met, Gabra1, Tnr, Glra2, Lama2, Gria1, Usp43, Rgs6, Mctp1, Rasgrf2, Cacna2d3, Sla, Cd200, Fbn2, Apba1, Prkar1b, Grin1, Dync1i1, Grip2, Grin2b, Ptpro, Slc6a11, Clstn2, Arpp21, Cck, Gucy1a3, N28178, Fam135b, Trim67, Dnm3, Cdh12, Necab1, Reps2, Rims1, Dlx1, 9330132A10Rik, Fam19a2, Ccbe1, Gpr85, Lingo1, Dlg2, Adap1, Fry, Ryr3, Camk2b, Unc5c, Kcnma1, mt-Tf, mt-Tv, mt-Nd4l, Nrxn3, A830039N20Rik, Scn2a1, Gm15344, Nwd2, Dlx6os1, Uba52, Gm26871, RP23-74F20.3, RP23-333L19.1, RP23-317L18.3, RP23-14P23.8, RP23-14P23.9 |

(d) **Covering markers of Telley dataset with $d = 15$ and $\alpha = 0.02$ extended from markers in Table 4c**

| | |
|---|---|
| 1H | Nr2e1, Zfp36l1, Gli3, Spata13, Ajuba, Col2a1, Hes1, Flnb, Notch2, Cenpa, Adamts9, Dusp16, Tead2, Sall1, Gsta4, Aspm, Etl4, Kif15, Ildr2, Rbm20, Gm12481, Sox3, Cenpe, Cldn12, Gas1, Creb5, Cdca7, Dach1, Tcf7l1, Hmga2, Gm7536, Rps18-ps3, Zfp516, Gm8129, Rps26-ps1, Rps10-ps1, Rpl15-ps3, Sox21, Phactr2, Rpl34-ps1, Rps13-ps2, Sox2, Gm10224, Gm6654, Gm3362, Gm13267, Gm15795, Hspa1b, Hspa1a, Gm26870, Gm28625, WI1-1003N17.1 |
| 24H | Rnd2, Kcnn1, Kit, Syn2, Myt1, Dusp14, Nuak1, Gadd45g, Lrrc16b, Wnt7b, 5330426P16Rik, Serping1, Tmem178, Rcor2, Ifitm3, Nrp1, Pramel7, Tuba4a, Slc30a10, Sema6d, Necab3, Rasgef1b, Cabp1, Tmem176b, Nkd1, Sh3bgrl2, Crabp1, Eomes, Shf, Neurod1, Klhl35, Abca7, Ppp1r14a, Osbpl5, Inf2, Ttc39b, Clvs1, Sox5, Pxylp1, Shb, Rpsa-ps10, Sstr2, Bcl11b, Nhlh2, Plcb1, Nhlh1, 9630028B13Rik, Rpl31-ps13, Cntn2, A930024E05Rik, Itpk1, Scrt2, Cxcl12, Unc5d, mt-Ti, mt-Co3, Adamtsl3, Gm23935, Gm4294, Gm12005, Gm15481, Rpl38-ps1, Gm11989, Gm14094, Gm15232, Gm13789, Pcp4, Gm6467, Rps19-ps12, A730020E08Rik, Gm5619 |
| 96H | Gabra2, Cacna1e, Chd5, Ndrg1, Mef2c, Gabrb2, Crlf1, Met, Gabra1, Tnr, Cacna1d, Glra2, Lama2, Gria1, Usp43, Rgs6, Mctp1, Rasgrf2, Cacna2d3, Sla, Cd200, Fbn2, Apba1, Prkar1b, Atp1b1, Grin1, Sparcl1, Dync1i1, Npy, Grip2, Grin2b, Ptpro, Slc6a11, Clstn2, Cspg5, Arpp21, Cck, Gucy1a3, Syt1, N28178, Fam135b, Trim67, Gabbr2, Dnm3, Cdh12, Necab1, Reps2, Inhba, 4930506M07Rik, Rims1, Dlx1, 9330132A10Rik, Fam19a2, Ccbe1, Gpr85, Lonrf2, Lingo1, Dlg2, Arhgap20, Adap1, Fry, Ryr3, Camk2b, Unc5c, Gm10115, Kcnma1, mt-Tf, mt-Tv, mt-Tl1, mt-Nd4l, Nrxn3, A830039N20Rik, 1810009A15Rik, Scn2a1, Gm15344, Gm15662, Nwd2, Dlx6os1, Uba52, Gm26871, RP23-74F20.3, RP23-333L19.1, RP23-317L18.3, RP23-14P23.8, RP23-14P23.9 |

(e) **Covering markers of Telley dataset with $d = 20$ and $\alpha = 0.02$ extended from markers in Table 4d**

| | |
|---|---|
| 1H | Otx1, Cdc20, Nr2e1, Plagl1, Grb10, Zfp36l1, Gli3, Samd4, Spata13, Ajuba, Col2a1, Hes1, Flnb, Notch1, Wwtr1, Notch2, Cenpa, Adamts9, Dusp16, Tead2, Sall1, Gsta4, Aspm, Etl4, Kif15, Mavs, Ildr2, Zic5, Dmrt3, Rbm20, Gm12481, Sox3, Cenpe, Cldn12, Gas1, Creb5, E330013P04Rik, Fgfr3, Cdca7, Dach1, Tcf7l1, Hmga2, Gm7536, Rps18-ps3, Zfp516, Gm8129, Rps26-ps1, Rps10-ps1, Rpl15-ps3, Sox21, Gm5453, Phactr2, Rpl34-ps1, Rps13-ps2, Sox2, Gm10224, Gm6654, Gm3362, Gm13267, Gm15795, Hspa1b, Hspa1a, Gm26870, 1190002F15Rik, Gm28625, WI1-1003N17.1 |
| 24H | Sez6, Rnd2, Kcnn1, Kit, Syn2, Myt1, Dusp14, Nuak1, Chga, Gadd45g, Lrrc16b, Wnt7b, 5330426P16Rik, Serping1, Tmem178, Rcor2, Ifitm3, Nrp1, Pramel7, Tuba4a, Slc30a10, Sema6d, Necab3, Trp53inp1, Rasgef1b, Cabp1, Tmem176b, Nkd1, Sh3bgrl2, Crabp1, Eomes, Shf, Neurod1, Klhl35, Abca7, Ppp1r14a, Osbpl5, Inf2, Neurod6, Ttc39b, Neurod2, Nrn1, Ccser1, Clvs1, Sox5, Kif21b, Mfap4, Pxylp1, Shb, Rpsa-ps10, Sstr2, Bcl11b, Nhlh2, Plcb1, Nhlh1, 9630028B13Rik, Rpl31-ps13, Cntn2, A930024E05Rik, Itpk1, Rpl31-ps20, Scrt2, Cxcl12, Unc5d, mt-Ti, mt-Co3, Tecpr1, Hist2h2ac, Adamtsl3, Gm23935, Gm4294, Gm12005, Gm15481, Rpl38-ps1, Gm8606, Gm11989, Gm14094, Gm15232, Gm13789, Pcp4, Gm6467, Rps19-ps12, A730020E08Rik, Gm5619, 2600014E21Rik, Gm8885 |
| 96H | Gabra2, Dlgap1, Cacna1e, Chd5, Ndrg1, Mef2c, Runx1t1, Gabrb2, Crlf1, Met, Gabra1, Prrt1, Tnr, Cacna1d, Glra2, Cacng2, Lama2, Gria1, Usp43, Rgs6, Mctp1, Rasgrf2, Cacna2d3, Sla, Cd200, Grm2, Nrxn1, Fbn2, Apba1, Prkar1b, Atp1b1, Grin1, Ntsr1, Ppp2r2c, Sparcl1, Dync1i1, Npy, Grip2, Grin2b, Ptpro, Tmtc1, Slc6a11, Clstn2, Cspg5, Arpp21, Cck, Gucy1a3, Lrfn5, Syt1, N28178, Fam135b, Trim67, March4, Gabbr2, Dnm3, Cdh12, Necab1, Reps2, Inhba, 4930506M07Rik, Rims1, Dlx1, Bend6, 9330132A10Rik, Fam19a2, Syt16, Amer3, Nxph1, Ccbe1, Gpr85, Lonrf2, Lingo1, Grm5, Dlg2, Arhgap20, Ppfia2, Cntn1, Maf, Adap1, Fry, Ryr3, Camk2b, Unc5c, Erbb4, Opcml, Gm10115, Kcnma1, mt-Tf, mt-Tv, mt-Tl1, mt-Nd4l, Nrxn3, Gad1, A830039N20Rik, 1810009A15Rik, Scn2a1, Gm15344, Gm15662, Nwd2, Dlx6os1, Uba52, A330076H08Rik, Gm26871, RP23-74F20.3, RP24-204F16.4, RP23-416H10.5, RP23-333L19.1, RP23-317L18.3, RP23-14P23.8, RP23-14P23.9 |

(f) **Covering markers of Telley dataset with $d = 25$ and $\alpha = 0.02$ extended from markers in Table 4e**

**Supplementary Table 4: CellCover marker genes extension at various depth.**

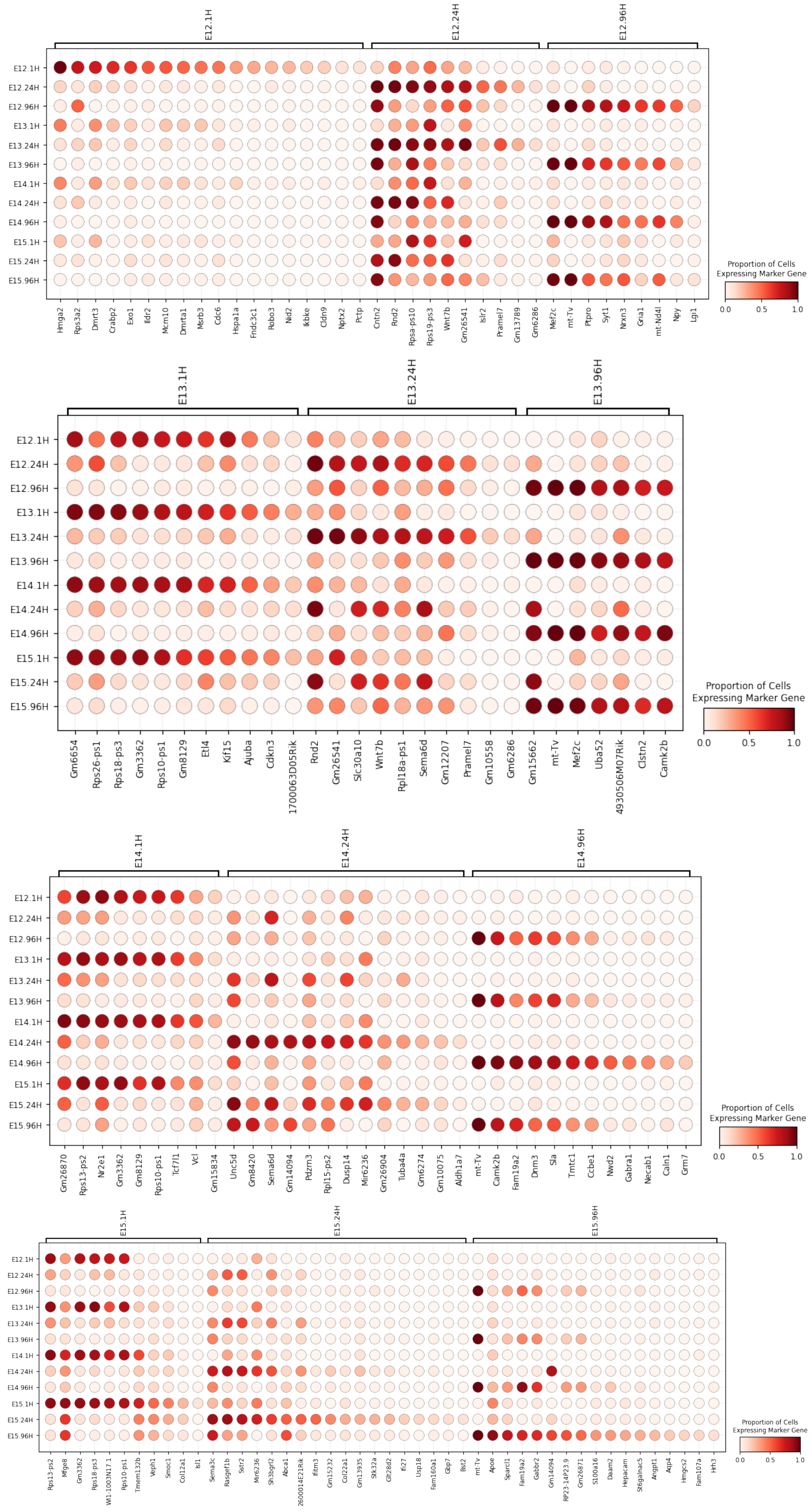


**Supplementary Figure 6**: **Dot plot of CellCover marker gene panels of three cell ages (1, 24, and 96hr) aggregated across embryonic days (E12 - 15) in mouse neocortical development.** The marker genes of each cell age at its corresponding embryonic day are grouped along the horizontal axis and sorted by expression frequency in the cell class of interest. The color of the dots represents the expression probability of covering markers conditioned on time since a cell's terminal division. The panel is obtained using the CellCover with $\alpha = 0.02$ and $d = 5$.

### 5 Genes in covering panels and marker genes defined by differential expression (DE) capture distinct elements of cell-type specific gene expression

Currently, the vast majority of cell type markers derived from scRNA-seq data are selected from the top of a ranked list of genes differentially expressed between a cell class of interest versus other cell types [7, 8]. This approach has yielded many insightful results across tissue systems and species. While this method optimizes individual genes as cell type markers (by their DE rank) and creates a set of markers by selecting top individual genes that distinguish a cell type, CellCover optimizes a panel that functions together to define a cell class. This fundamental difference in the search for marker genes results in the capture of distinct cell-type specific signals. Therefore we put forth CellCover as a complementary approach to conventional DE marker gene finding, rather than a competing method for cell-type marker gene discovery.

Notable among the additional differences in these marker-finding methods is that CellCover (1) obviates the need for normalization, scaling or feature selection (as a function of utilizing binarized expression data), and (2) can directly capture heterogeneity within a cell class of interest. We show the latter empirically by comparing markers derived from CellCover and DE methods in the Telley data. Figure S7 summarizes the overlap of covering marker gene panels in Figure S6 (NeMO: Telley 12 CellCover Panels) and the Seurat DE ranked genes (NeMO: Telley 12 Sets DE Genes) for all 12 cell classes. In the majority of cell classes, there is appreciable divergence in the marker genes that the two methods identify as cell-type specific (represented by vertical gaps between the red and blue lines in each plot). Concordance between 96H cell markers is greatest (perhaps indicating lower within-class heterogeneity in neurons), with the shortest list of top DE selected markers that include an entire covering marker gene panel at 40 genes in length (E13.96H). These observations indicate that in practice, the two methods of marker gene discovery produce quite divergent results.

To further explore the discordance in marker genes selected by the two methods, we visualized expression levels of individual E12.1H covering markers with high p-value rank in the DE (Figure S8 top panel) and those that are in the CellCover panel but rank lowly in the DE ranked list (Figure S8 bottom panel). We find markers with high DE rank, e.g., Hmga2 and Crabp2, are predominantly expressed in E12.1H cells, and they are also expressed in other cell populations, though less frequently. In contrast, genes appearing in the CellCover solution but at low rank in the DE are less frequently expressed but are more specific to the E12.1H class. For example, covering marker genes Nptx2, Pctp2, and Clnd9 are not as frequently expressed in E12.1H cells as other markers. However, these genes are rarely expressed in other cell classes. These marker genes with high specificity are very discriminative for the purpose of defining cell classes when used together but may be omitted by DE marker selection methods in the feature selection or DE calculation steps due to low and/or less frequent expression, i.e., low sensitivity as an individual marker. By optimizing the panel of marker genes rather than individual markers, CellCover can borrow power across multiple genes that show greater specificity but which individually do not identify all cells in a class.

To show this more systematically, we ran differential expression analysis on the 1H, 24H, and 96H cells in the Telley data [4] and showed the rank of the CellCover markers (Figure 2A in the main manuscript) based on the p-values returned from the DE analysis in Figure S9. We found that across all the cell ages, the CellCover markers that have low sensitivity also have a lower rank in the DE. For example, in the 1H, the CellCover marker Hspa1a has the lowest sensitivity among all markers, expressed only near 13% percent of the time in this cell age, and its rank from DE is more than 500 (Table S5). Although this gene has low sensitivity, it has high specificity, rendering it a good candidate for a marker gene when used with other complementary genes. Genes with these properties appear at much lower ranks in DE and are hence not selected when using that method.

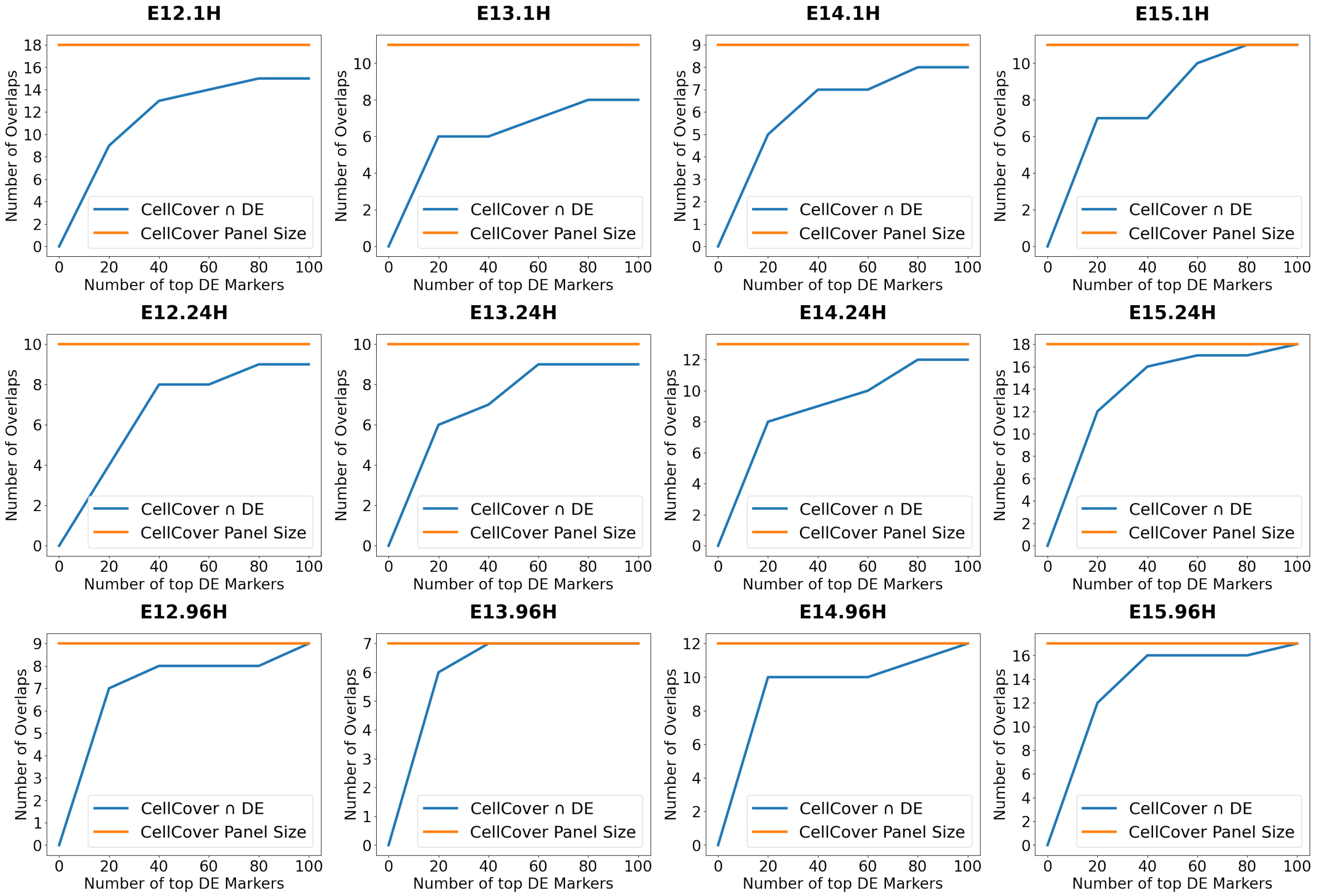


**Supplementary Figure 7**: **Overlap and discordance between marker genes selected by CellCover and DE methods.** For each of the 12 CellCover gene panels defined in the Telley mouse data, the number of overlapping genes found by DE methods is quantified. In each panel, the orange line indicates the number of genes in the CellCover gene panel for that cell class. The blue line indicates the cumulative number of CellCover genes also found in the top $x$ ranked DE gene marker list as increasingly large DE gene lists are considered. The X-axis value where the orange and blue lines meet indicates the number of DE markers that would need to be selected to include all the CellCover markers.

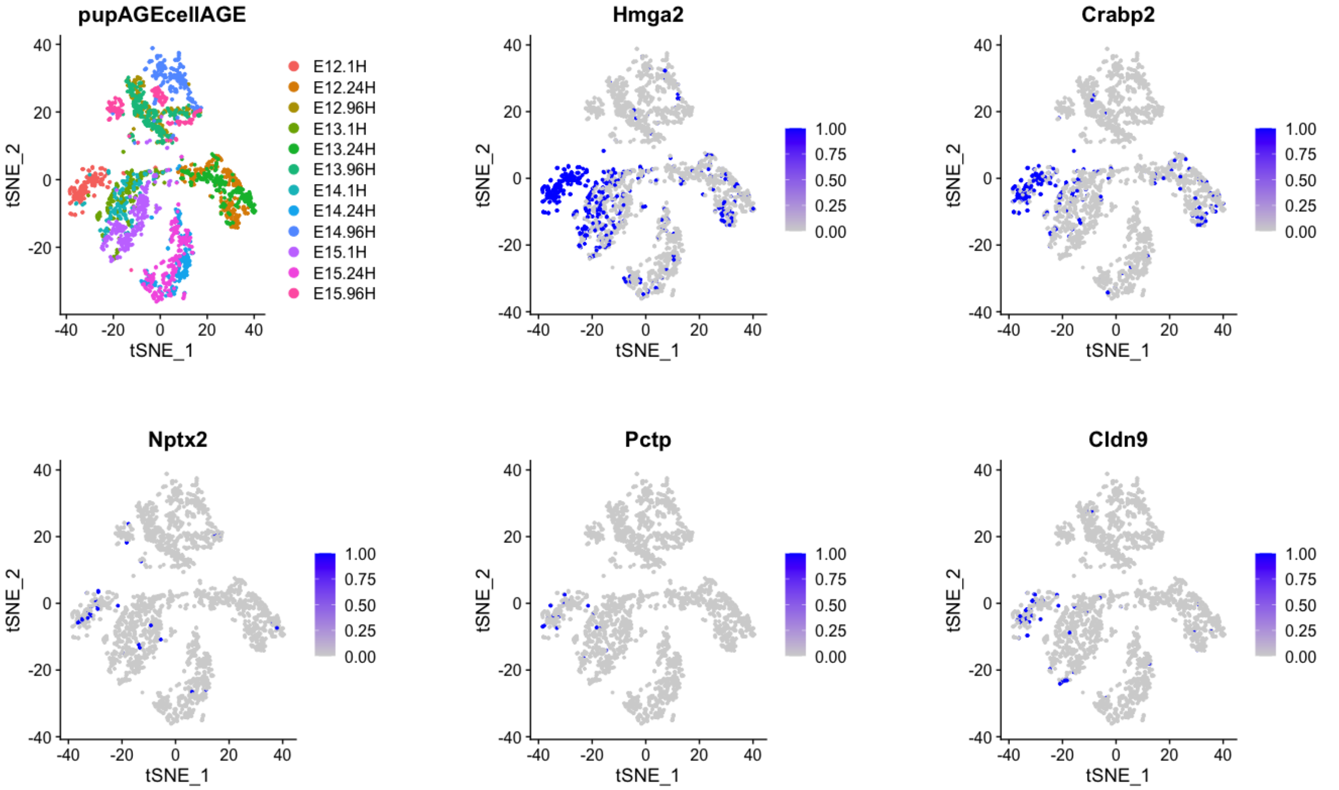


**Supplementary Figure 8**: **Sensitivity and Specificity of Individual marker genes selected by CellCover:** Visualization of expression of individual marker genes selected by CellCover. Among all the covering markers for E12.1H, Hmga2 and Crabp2 have the lowest p-values in DE (and are therefore selected as DE marker genes) and show high sensitivity with lower specificity. In contrast, Nptx2, Pctp, and Cldn9 have the highest p-values in DE (and are therefore not selected as DE marker genes) and show low sensitivity individually, but very high specificity. This ability to combine highly specific genes with individually low sensitivity - to generate marker panels of both high specificity and sensitivity - is responsible for much of the differences between gene panels selected by CellCover versus DE methods.

#### 6 Distinction between CellCover and PhenotypeCover

In addressing the challenge of marker gene selection, another approach, PhenotypeCover [9], also draws inspiration from the set covering problem. While PhenotypeCover deploys multiset multicover to identify a global set of marker genes that discriminate all the cell types, CellCover captures cell-type specific signals whose combinatorial patterns characterize the gene expression behavior of the target cell sub-population, with the flexibility to extend to a global covering. A key distinction between the two methods lies in the resolution of the covering. CellCover defines covering at the level of both individual cells and cell-types, in contrast to the global cover of the entire cell population defined in the PhenotypeCover. This enables CellCover to better explore cellular heterogeneity and inspect the dynamics of different cell types in particular biological processes. Moreover, the difference in the weighting scheme of the two methods gives rise to the different properties of marker genes. Like the comparison with DE, our results in Figure S10 show that PhenotypeCover favors discriminative genes that are highly expressed. In contrast, CellCover selects genes from a wider spectrum regarding the expression rate, where highly specific, rarely expressed genes are often selected. Important mathematical distinctions between the two methods are also addressed in the following paragraphs.

The object to be covered are different in the two methods. CellCover defines covering at level of both individual cell and cell-type. We say a cell is covered by marker panel $Q$ if more than $d$ genes in $Q$ are expressed in the cell. What is more, a cell type $k$ is covered if more than $1-\alpha$ proportion of the cells in this cell-type are covered. CellCover then identifies the optimal marker panel by minimizing the sum of weights of genes in the marker panel, where the weight of each gene reflects the inverse of its differentiating power of cell-type $k$, subject to the covering constraints on this cell-type. Therefore, Cellcover identifies a marker panel for each cell-type such that the marker

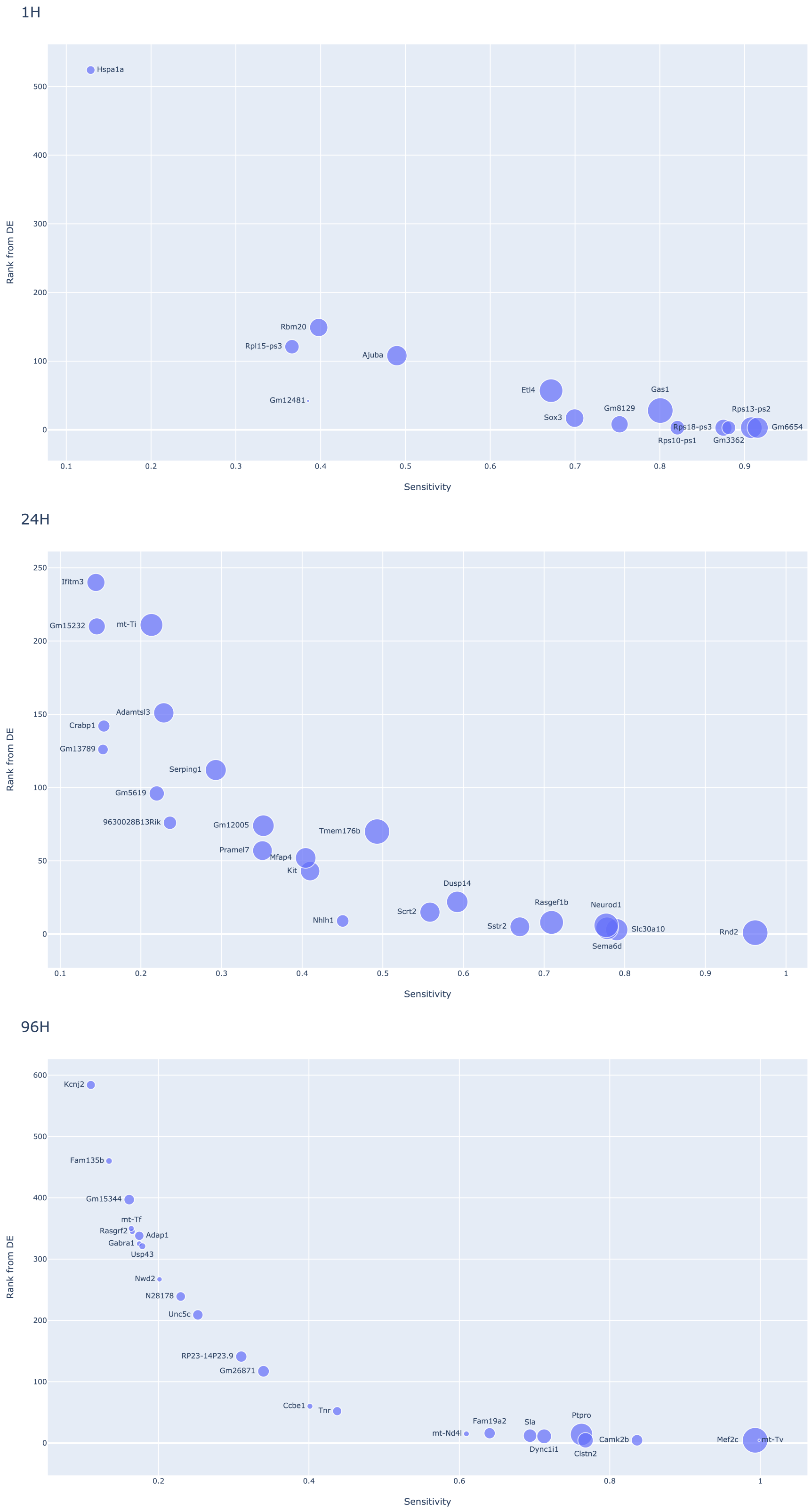


**Supplementary Figure 9**: **CellCover markers sensitivity vs rank from DE.** We ran Seurat DE on the Telley dataset to discriminate 1H, 24H, and 96H and rank the entire gene portfolio by the p-values. Each CellCover gene $g$ (from Figure 2A in the main paper) has its sensitivity of the class $k$, $P(X_g > 0|Y = k)$, on the horizontal axis and its rank from DE on the vertical axis. The size of the dot represents the weight of the marker defined in the method section (download the interactive plots at DE Rank vs Sensitivity). The ability of CellCover to borrow power across marker genes of low sensitivity and high specificity is one of the primary factors that sets it apart from DE methods.

| Gene | $\frac{\sum_{l\neq k} P(U_g=1\|Y=l)}{K-1}$ | $P(U_g=1\|Y=k)$ | Rank in DE |
|---|---|---|---|
| Rps10-ps1 | 0.07 | 0.82 | 3 |
| Gm3362 | 0.07 | 0.88 | 3 |
| Gm6654 | 0.17 | 0.92 | 3 |
| Rps18-ps3 | 0.10 | 0.88 | 3 |
| Rps13-ps2 | 0.17 | 0.91 | 3 |
| Gm8129 | 0.09 | 0.75 | 8 |
| Sox3 | 0.10 | 0.70 | 17 |
| Gas1 | 0.21 | 0.80 | 28 |
| Gm12481 | 0.00 | 0.39 | 42 |
| Etl4 | 0.15 | 0.67 | 57 |
| Ajuba | 0.08 | 0.49 | 108 |
| Rpl15-ps3 | 0.03 | 0.37 | 121 |
| Rbm20 | 0.05 | 0.40 | 149 |
| Hspa1a | 0.00 | 0.13 | 524 |

**Supplementary Table 5**: **Sensitivity analysis of CellCover 1H markers.** The first column of the table lists the average sensitivity of the marker gene in the cell population that is not 1H, and the second column lists the sensitivity of the marker within class 1H. The third column lists the p-value rank from DE. The CellCover marker with low sensitivity of 1H also tends to have a low p-value rank in DE.

genes not only possess discriminative power of the phenotype but also cover the phenotype. Instead, PhenotypeCover identifies a global marker panel for the entire cell population such that the marker panel is of minimal cardinality and separates the phenotypes. In particular, in PhenotypeCover, a marker panel $Q$ provides a cover for the dataset if, for every phenotype pair $(i, j)$, the sum of weights of each gene in the marker panel exceeds $B$, where this weight of the gene depends on $(i, j)$ and are positively correlated with its discriminative power of this phenotype pair, and $B$ is a user-specified covering threshold. We note that CellCover also has a natural extension to identify a global marker set. To make a fair comparison, we will use the global version of the CellCover on the log normalized count data.

Let $N$ be the total number of cells and $G$ denotes the set of genes. Given the gene expression matrix $X \in \mathbb{R}^{N\times|G|}$ and a known vector y of length N representing the phenotype. We derive $M_{i,g}$ from X by averaging the log normalized expression of gene g across all the cells with phenotype $i$. We also derive $U \in \{0,1\}^{N\times|G|}$ through binarization, where we let $u_g^{(c)} = 1$ if $x_g^{(c)} > 0$ and 0 otherwise, and we say a gene is expressed if $u_g^{(c)} = 1$. Further, we let $Z \in \{0,1\}^{|G|}$ to a binary vector so that $z_g = 1$ means gene $g$ is selected in the marker panel. $Z$ is the variable we want to solve for in both methods. Next, we compare the mathematical forms of the two methods, and we will address the key differences from them.

**CellCover (Global Version)**: In the CellCover, besides $Z$, we will also optimize another auxiliary binary variable $s^c$, where $s^{(c)} = 1$ means that cell $c$ is covered. We also introduce two interpretable hyperparameters:

- $\boldsymbol{d}$: the covering depth. It is also the lower bound of the cardinality of the optimal covering panel.
- $\boldsymbol{\alpha}$: one minus the desired covering rate.

In the global version of CellCover, we define the weight to be $w_g = \min_k w_{g,k}$ where $w_{g,k} = \frac{\sum_{l\neq k} M_{g,l}/(K-1)}{M_{g,k}}$ and $M_{g,k} = \frac{\sum_{c\in S_k} x_g^{(c)}}{|S_k|}$. This weight reflects the maximum capacity of gene $g$ in differentiating one cell type versus the others. The integer program is given by

$$\min_Z \sum_{g\in G} w_g z_g \tag{1}$$

$$\sum_{g\in G} z_g u_g^{(c)} \geq \boldsymbol{d} s^{(c)} \quad c = 1, 2, ...N \tag{2}$$

$$\sum_{i=1}^{N} s^{(c)} \geq (1-\boldsymbol{\alpha})N \tag{3}$$

**PhenotypeCover:** The hyperparameter in PhenotypeCover is $B$, a user-specified threshold of multiset multicover. The integer program is given by

$$\min_{Z} \sum_{g \in G} z_g \tag{4}$$

$$\sum_{g \in G} |M_{g,i} - M_{g,j}| z_g \geq B \quad \forall \text{ cell-type pairs } (i,j) \tag{5}$$

Next, we list some key difference between the two methods:

• Weight scheme: The choices of weight are different from the two methods. CellCover uses ratio of the average expressions to define the weight, where PhenotypeCover uses the absolute difference of the average expression as the weight. One immediate advantage of the the CellCover weighting scheme is that it will encourage the algorithm to select genes that are specific to a phenotype which may have low overall expression rate. This may introduce some "noisy" genes to the marker panel, but also help us to identify rare, lowly expressed genes that are equally important to characterize the cell-type. On the other hand, the Phenotype weighting scheme, together with its constraint, favors only those genes that are frequently expressed. The algorithm requires the sum of weights to exceed $B$ for all cell-type pairs. Including a lowly expressed but highly specific gene will not only make constraints in PhenotypeCover harder to satisfy but also incur a cost to the objective function. In contrast, CellCover encourages the identification of rare yet specific genes, and it also favors the genes that are frequently expressed and cell-type differentiating because the marker panel needs to satisfy the covering constraints. Hence, we should expect the covering to find a broader spectrum of marker genes regarding the rate of expression than the PhenotypeCover.

• Solver: CellCover solves the constraint optimization problem using integer programming (branch and bound algorithm), so the answer is (almost) exact when the primal objective and dual objective bounds converge. In particular, the returned solution is in some $\epsilon$ ball of the true optimal solution. PhenotypeCover solves the optimization by using the greedy approximation algorithm for multiset multicover, in which the accuracy of the solution is upper bounded by a factor of $H_m$ increase in the solution size where $H_m = 1+1/2+...+1/m$ and $m$ is the cardinality of the largest multiset. CellCover provides a more accurate solution, but branch and bound algorithm has a slower run time than the greedy algorithm used by PhenotypeCover, which almost linear in the number genes considered). However, we note that using greedy algorithm to solve for set covering problem inevitably undermine the combinatorial nature of the problem. When applying PhenotypeCover to real datasets, many returned solutions will fail to satisfy the constraints defined in Inequality 5.

• The $B$ in the PhenotypeCover less directly inform the cardinality of the marker panel, and this threshold may differ from dataset to dataset due to difference in expression level. Determining a proper threshold in relation to the desired size of the marker panel requires trials and errors. In contrast, the hyperparameter $d$ in CellCover more directly informs the size of the marker panel, as it provides a lower bound to the number of markers in the optimal panel. Also, it is not dataset dependent.

• PhenotypeCover does not require covering on individual cell level. Therefore, the method does not assess the interactions among the marker genes in each cell. CellCover imposes restriction on the collective expression behavior of marker genes in single-cell level. This constraint leverages the biological facts that co-expression of marker genes may activate certain functions.

We compare the two methods by applying them to obtain the global marker set of the Telley dataset. The goal is to obtain a set of marker genes for the entire dataset that distinguishes the time stamps 1H, 24H, and 96H. For CellCover, we let $\boldsymbol{d} = 20$ and $\boldsymbol{\alpha} = 0.02$ to obtain a marker panel of size 61. For the PhenotypeCover, we set $B = 95$ to get a marker set of the same size as CellCover. According to Supplementary Figure 10, in comparison to PhenotypeCover, CellCover markers exhibit a larger range of probability of expression, encompassing not only highly expressed and discriminative genes but also encouraging the selection of rarely expressed yet highly specific genes.

### 7 Run Time Analysis and Stopping Criterion

The runtime performance of CellCover exhibits variability across different cell populations. This variability is not solely attributable to the size of the cell population under analysis. In practice, we have observed that CellCover can sometimes execute faster on datasets with larger cell populations than those with smaller ones. In general, the size of the effective search space of the candidate genes, the level of transcriptional distinctness of the cell type, and the size of the target cell population may jointly affect the runtime of our algorithm. Despite the potential for prolonged

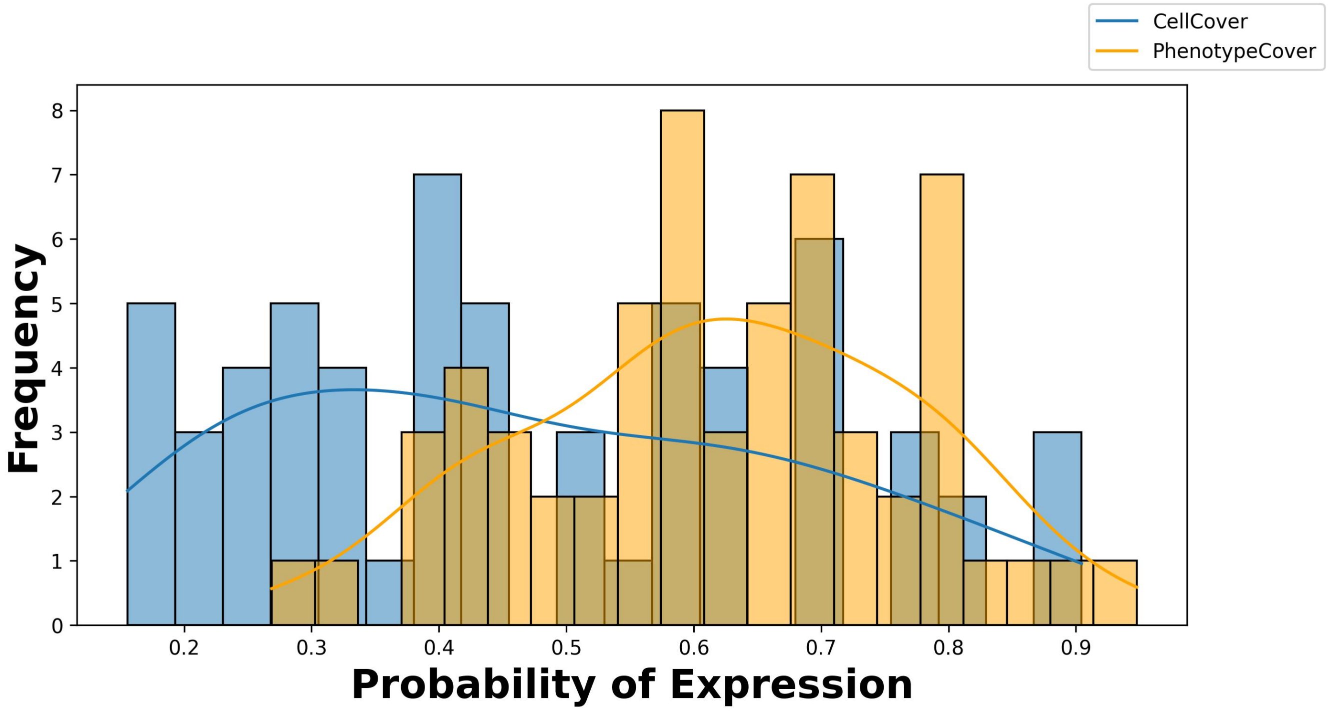


**Supplementary Figure 10**: **Distribution of marker gene expression rate of CellCover and PhenotypeCover**. The blue bars form the histogram of the probabilities of expression of the CellCover (global version) marker genes obtained at $\boldsymbol{d} = 20$ and $\boldsymbol{\alpha} = 0.02$. The probability of expression is computed by $\frac{\sum_{c=1}^{N} u_g^{(c)}}{N}$. The yellow bars depict the distribution of probabilities of expression of the Phenotype marker gene obtained by letting $B = 95$ to match the size of CellCover marker panel.

runtimes due to the NP-hardness nature of the branch and cut algorithm underlying the Gurobi solver, it is important to note that, once converged, CellCover guarantees a globally optimal solution. When the target cell population is large, even though we may not get a convergence certificate in a few seconds, the intermediate solution returned by Gurobi is guaranteed to be feasible (i.e., the corresponding gene panel will cover the target cell population) and can reach a stable suboptimal solution in a reasonable amount of time, typically within an hour in the worst case. A detailed analysis of the runtime is provided here, and we also outline a rule of thumb for determining the stopping criterion when the CellCover solver is slow on some cell population.

• Size of the target cell population: The number of the cells in a population of interest directly impacts the run time of covering. In CellCover, we define for each cell an auxiliary variable $s^{(c)}$ which indicates whether the cell $c$ is covered, and we will solve for this variable via integer programming. Hence, the number of the variables $s^{(c)}$ scale with the size of the cell subpopulation, leading to a potential increase in runtime.

• Size of the effective search space of candidate genes: In the CellCover, we execute the pruning step before inputting the data into the gurobi. Hence, the choice of filtering (See the Method section) hyperparameters affects the size of the candidate genes, which directly impacts the run time of CellCover. First of all, we only select genes of which the fraction of cells expressing it is above $t_e$. Hence, a small $t_e$ may slow down the run time since the integer program need to address the interplay of more genes. Secondly, for each cell type $k$, we select genes whose expression rates (the fraction of cells expressing the gene) within the subpopulation $k$ are higher than the average expression rates in other cell types. Equivalently, we filter out the genes with a margin less than the threshold $t_d$, for $t_d \geq 1$. In the implementation of the algorithm, since we might not know which $t_d$ is the desired cutoff, we execute this step of filtering by selecting the genes with the top $l$ margin in the target cell subpopulation that has a margin larger than 1. We observe that the requirement of $t_d \geq 1$ place the most significant filtering effect, sometimes reducing the number of candidate genes to the order of hundreds. As a result, the run time of CellCover scales with larger $l$ (or smaller $t_d$).

• The depth of covering $\boldsymbol{d}$: In general, a higher depth leads to an increase in run time because of the more difficult constraints to be satisfied. However, the impact of the covering depth also depends on the size of the candidate gene sets. If the covering depth is too high relative to the size of the effective search space of the genes, then the covering problem may become easier to solve, because the program will have to select a majority of the genes, if not all of them, to satisfy the constraints of covering. In this case, even though higher depth may surprisingly lead to a decrease in run time, the program becomes more susceptible to the feasibility problem because there may not be enough genes to cover the cell subpopulation at a large depth.

We augment this theoretical analysis with a run-time experiment on the Polioudakis dataset [5] to extract CellCover markers from each of the 10 cell types. These experiments are run on a desktop with Intel 13900K CPU, and the results are shown in Figure S11.

Next, in case when the CellCover may take long to converge, we add features in the CellCover package which will guide the user to monitor the convergence of the program and implement stopping criterion to obtain stable, suboptimal solution. When the CellCover starts running, it will output the log in real time. In the log, the metric "MIPGap" will be the convergence indicator. By definition, it is the gap between the lower (dual) and upper objective bound (primal). The MIP solver will terminate with an optimal result when the gap between the MIPGap is sufficiently small. By default, the terminating MIPGap threshold is $10^{-4}$. To implement early stopping in the simplest way, we can either

- increase the gap threshold by setting the parameter argument "MIPGapThreshold" in the python CellCover function to be any user-defined value.
- set a hard stopping time limit of the program by setting the parameter "TimeLimit" (in seconds).

We note that every intermediate solution returned by the integer programming is guaranteed to be feasible, meaning that even the primal-dual gap on the objective is non-zero, the current gene panel solution ensures that the covering constraints are satisfied.

We also include additional features to the program that aid the users in monitoring the convergence and stableness of the solution in real time, and help the user to terminate the program with confidence. In particular, whenever the Gurobi improves its bound, we will periodically ask the program to print the current genes panel being found which is the best solution so far. Each time the program makes a callback, we also compute the size of the set difference between the current solution and the last solution. If the solution is converging, then this set difference will ideally become smaller. We let the program terminate if the set difference mentioned above is less than 3 for more than three consecutive callbacks. Users can also terminate the program themselves and obtain the most recent solution at their convenience.

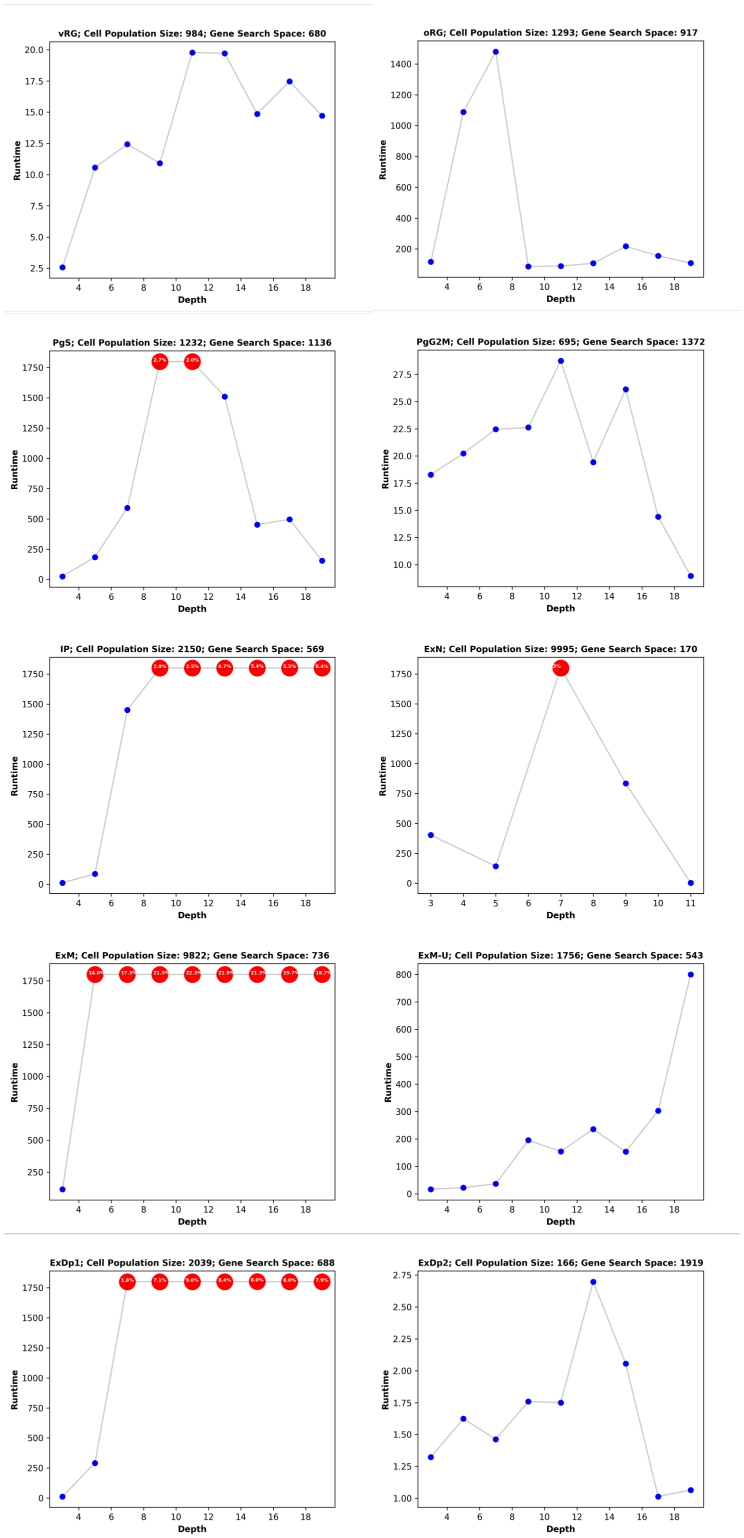


**Supplementary Figure 11**: **Covering run time of multiple cell populations from the Polioudakis dataset:** The $x$ axis is the covering depth $\boldsymbol{d}$, and the $y$ axis is the run time. We set a time limit of 1800 seconds for the program. If the branch and cut algorithm does not converge in 1800 seconds, MIP gap will be reported. The smaller the gap, the closer the dual and primal objectives are, indicating better convergence.

### 8 Additional Transfer of Covering Marker Gene Panels

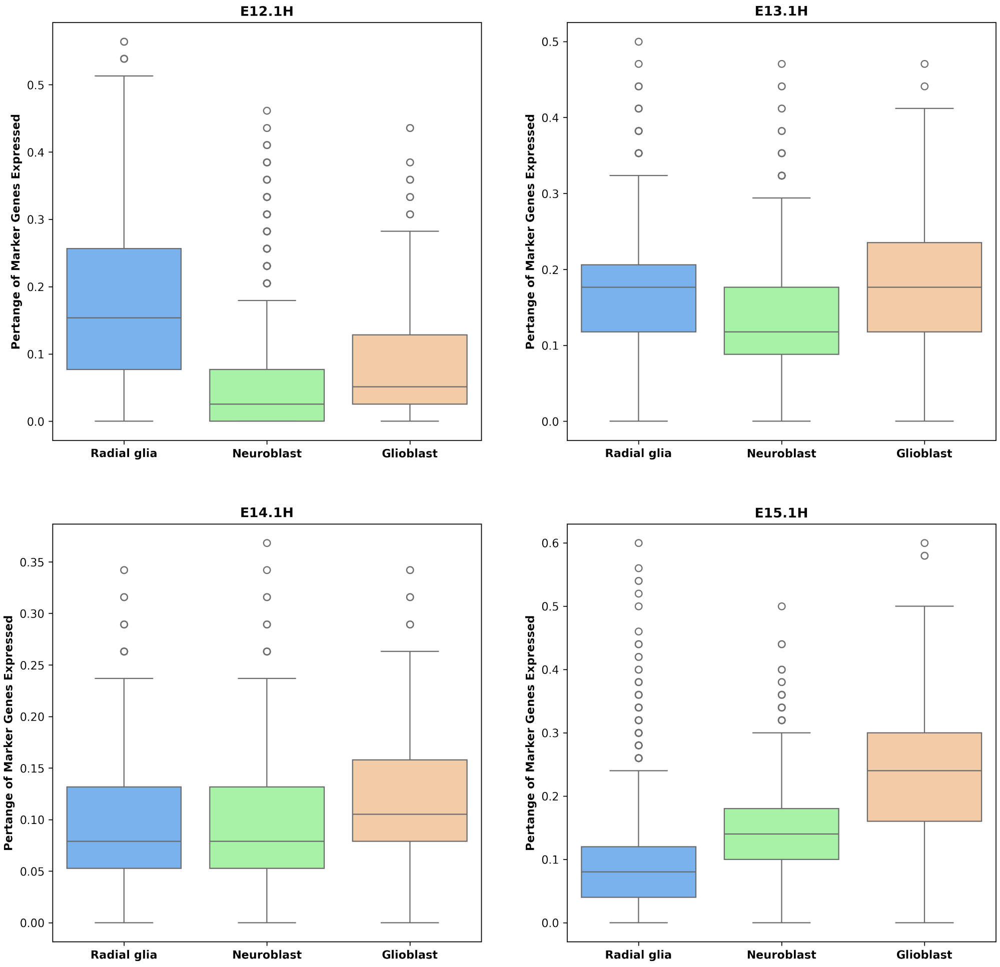


**Supplementary Figure 12**: **Transfer of Telley within 1H markers to radial glia, neuroblast, and glioblast cell populations in LaManno Dataset [10]:** We use the box plot to show the distribution of the proportion of marker genes expressed in each of the three cell population. For example, the first (blue) box plot in the top left panel shows the distribution of the proportion of E12.1H marker genes expressed in the radial glia cells in the LaManno dataset. These data clearly show the shift of mouse neocortical progenitors from to neurogenic to gliogenic potential from E12-E15.

### 9 Additional Cell Type Mapping Across Datasets Through Marker Transfer

To demonstrate that CellCover effectively captures cell-type-specific information across diverse single-cell RNA-seq datasets, we performed cross-dataset marker transfer experiments. Using Hao PBMC CITE-seq dataset [8] (Figure S13E) as source dataset with marker panels generated at depth 5 and a covering rate of 98%, we evaluated the transferability of CellCover markers to four target datasets (Figure S13A-D): Tabula Sapiens blood data [11] sequenced with three protocols (smart-seq2, 10x 3', and 10x 5') and the Stephenson COVID-19 PBMC dataset [12], restricted to cells from healthy controls.

For each transfer experiment, we mapped cell type $k$ in the source dataset to cell type $k'$ in the target dataset by calculating the proportion of $k'$ cells expressing at least five markers from the source marker panel $M_k$. Supplementary Figures S14A-D illustrate the results of these transfer experiments. The heatmaps reveal that the marker genes identified by CellCover are primarily expressed in the corresponding cell types in the target datasets, underscoring the robustness and cell-type specificity of the method across different sequencing technologies and biological contexts.

To augment the transfer results at cellular resolution, we also show in Figures S18-21 the proportion of genes in the CellCover marker panel $M_k$ expressed in each cell from the target dataset for all cell types $k$ in the source data. These figures highlight that the cells in the target datasets with the highest abundance of $M_k$ expression corresponds to cell states closely related to the source cell type $k$, further supporting the specificity and transferability of CellCover marker panels.

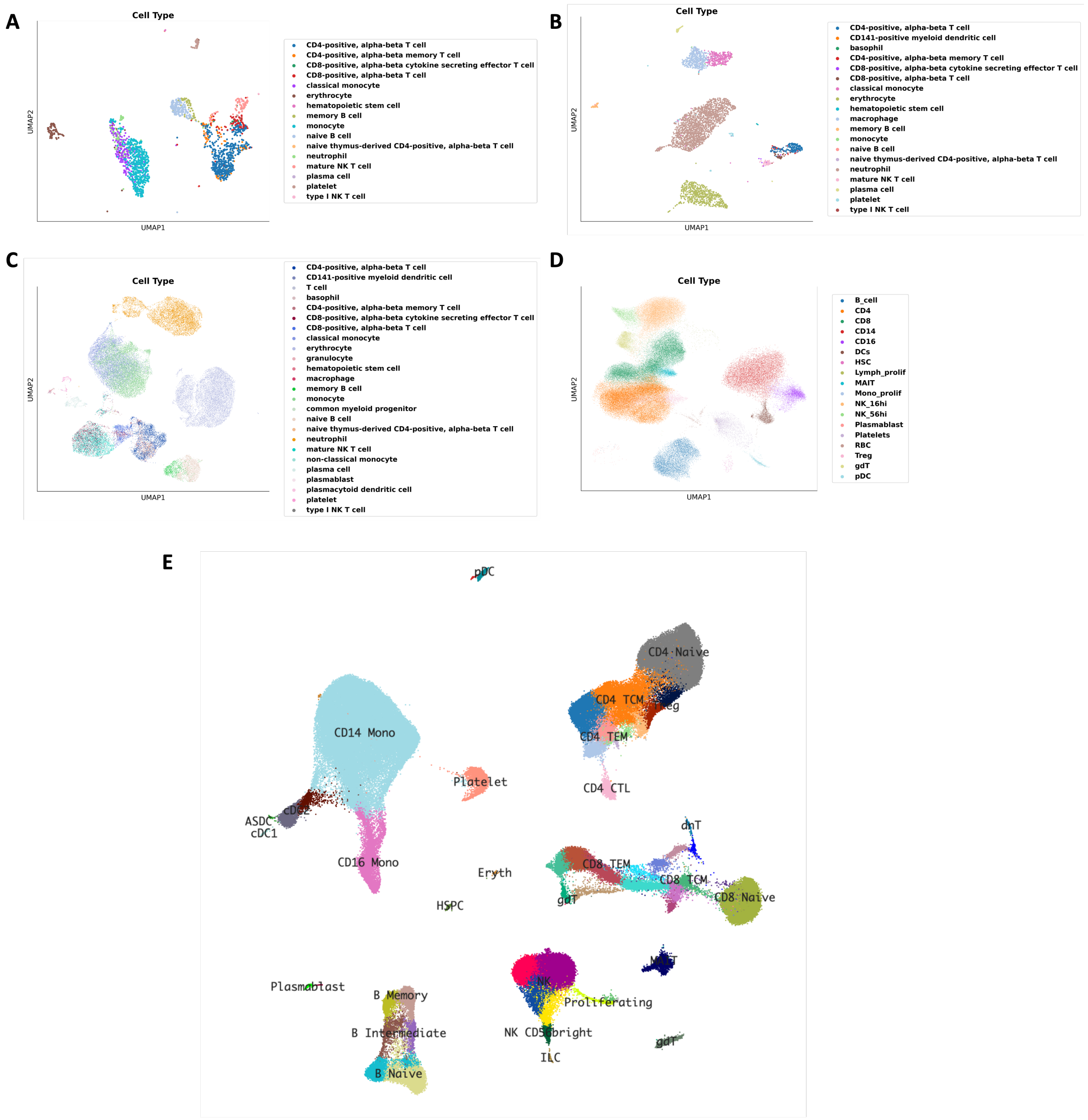


**Supplementary Figure 13**: **UMAP visualizations of cell type annotations for the Tabula Sapiens blood datasets, the Stephenson COVID-19 PBMC dataset and the Hao PBMC dataset.** A) Tabula Sapiens blood dataset sequenced using the Smart-seq2 protocol. B) Tabula Sapiens blood dataset sequenced using the 10x Genomics 5' protocol. C) Tabula Sapiens blood dataset sequenced using the 10x Genomics 3' protocol. D) Stephenson COVID-19 PBMC dataset. E) Hao PBMC Dataset where the UMAP visualization is available at https://atlas.fredhutch.org/nygc/multimodal-pbmc/.

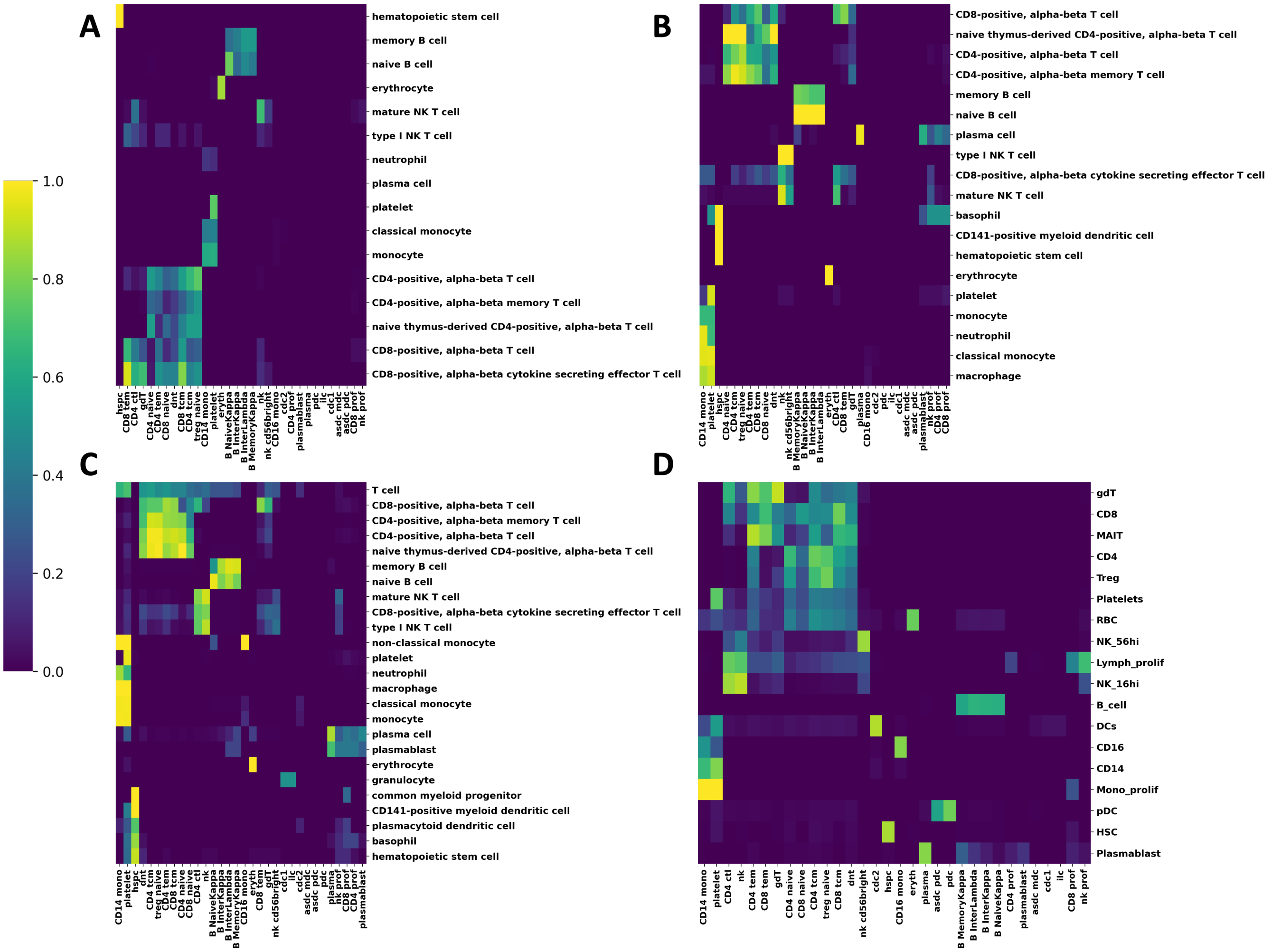


**Supplementary Figure 14**: **Cross-dataset marker gene transfer results.** Heatmaps show the proportion of target cells (y-axis) expressing at least five markers from source cell-type-specific marker panels (x-axis) generated by CellCover. Subplots (A–D) represent transfer results to Tabula Sapiens blood datasets sequenced with smart-seq2 (A), 10x 5' (B), 10x 3' (C), and the Stephenson COVID-19 PBMC dataset with only the healthy controls (D).

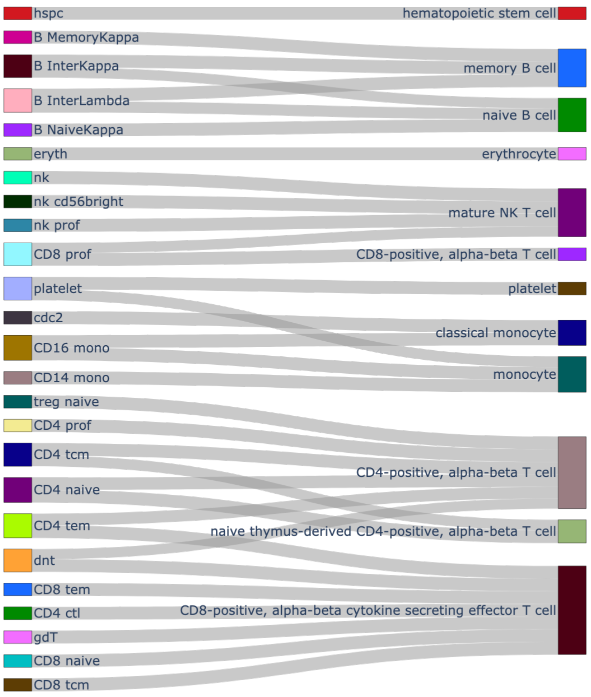


**Supplementary Figure 15: Sankey diagram depicting cell type mapping from the Hao PBMC CITE-seq dataset to the Tabula Sapiens blood dataset (Smart-seq Protocol)**

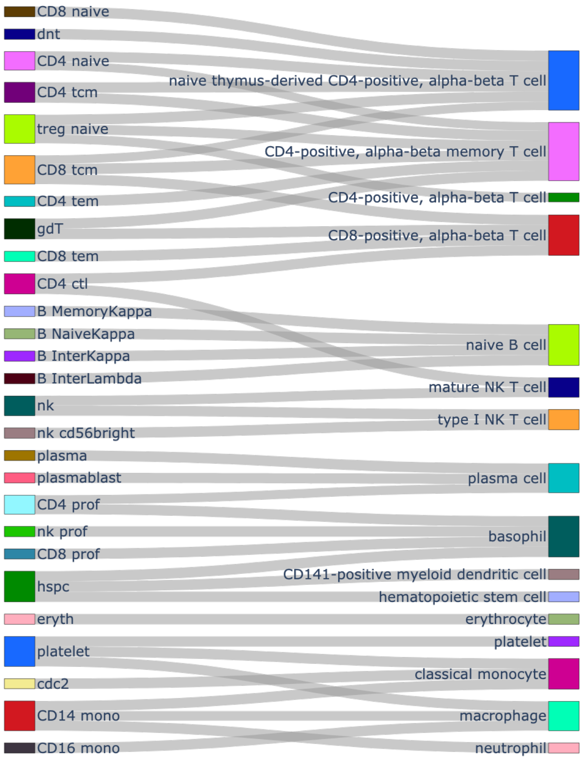


**Supplementary Figure 16: Sankey diagram depicting cell type mapping from the Hao PBMC CITE-seq dataset to the Tabula Sapiens blood Dataset (10x 5' Protocol)**

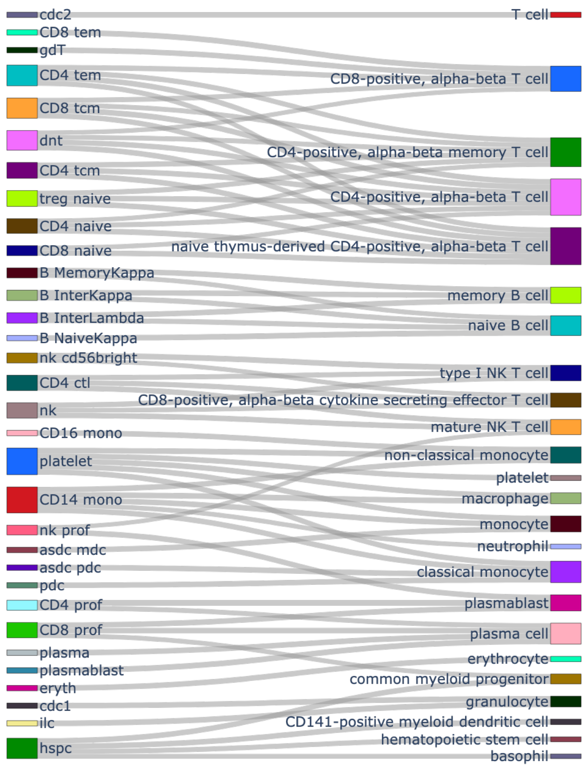


**Supplementary Figure 17**: **Sankey diagram depicting cell type mapping from the Hao PBMC CITE-seq dataset to the Tabula Sapiens blood dataset (10x 3' Protocol)**

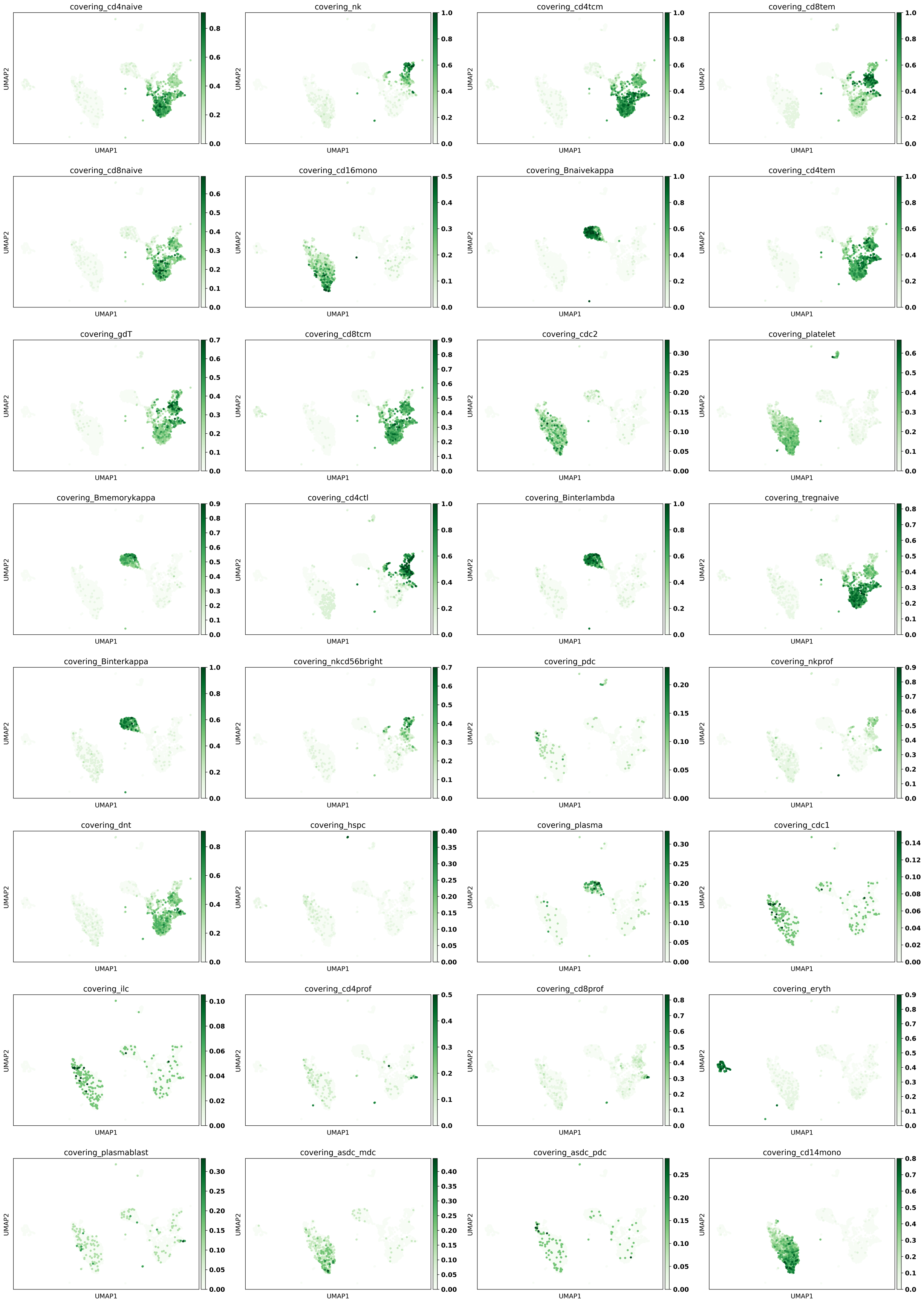


**Supplementary Figure 18**: **Transfer of CellCover marker panels from PBMC CITE-seq Dataset to Tabula Sapiens blood dataset sequenced by smart-seq2 protocol.** For each of the 32 cell types in the PBMC datasets, a CellCover marker panel of depth 5 and covering rate of 98% was derived. Each panel was transferred to the target dataset, with the plots showing the proportion of marker genes expressed in cells of the target dataset, as indicated by the titles of each subplot.

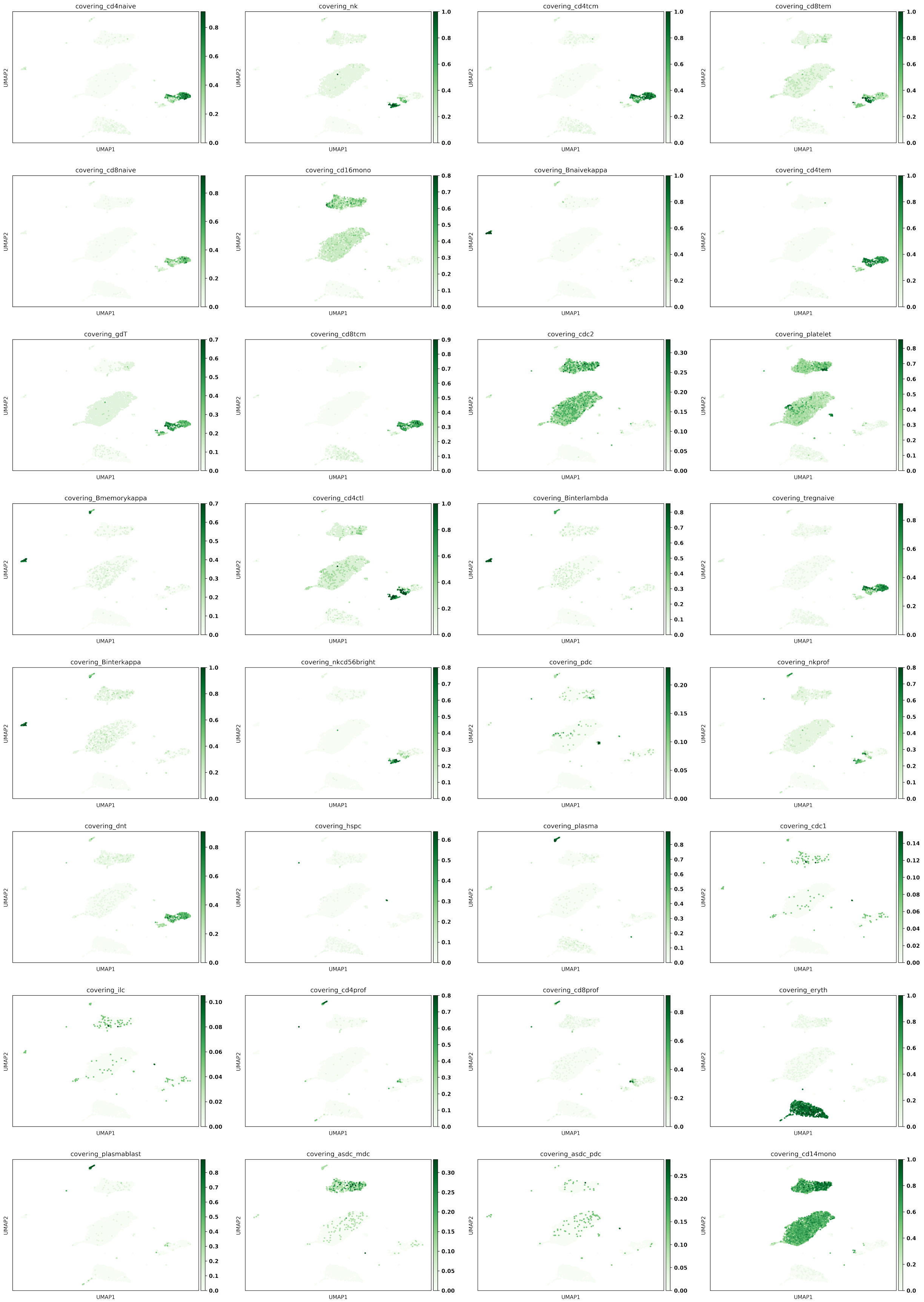


**Supplementary Figure 19**: **Transfer of CellCover marker panels from PBMC CITE-seq dataset to Tabula Sapiens blood dataset sequenced by 10x Genomics 5' protocol.** For each of the 32 cell types in the PBMC datasets, a CellCover marker panel of depth 5 and covering rate of 98% was derived. Each panel was transferred to the target dataset, with the plots showing the proportion of marker genes expressed in cells of the target dataset, as indicated by the titles of each subplot.

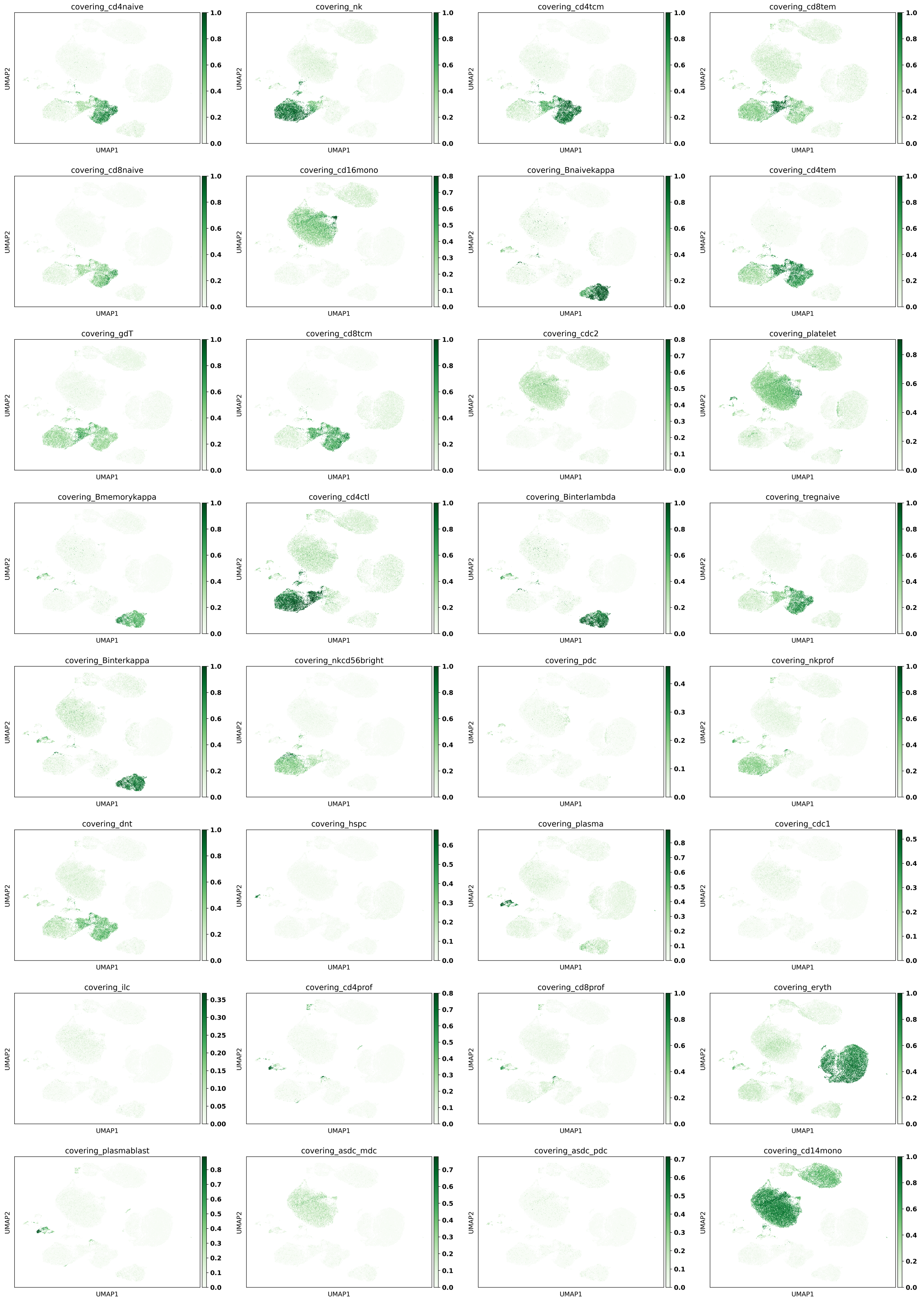


**Supplementary Figure 20**: **Transfer of CellCover marker panels from PBMC CITE-seq dataset to Tabula Sapiens blood dataset sequenced by 10x Genomics 3' protocol.** For each of the 32 cell types in the PBMC datasets, a CellCover marker panel of depth 5 and covering rate of 98% was derived. Each panel was transferred to the target dataset, with the plots showing the proportion of marker genes expressed in cells of the target dataset, as indicated by the titles of each subplot.

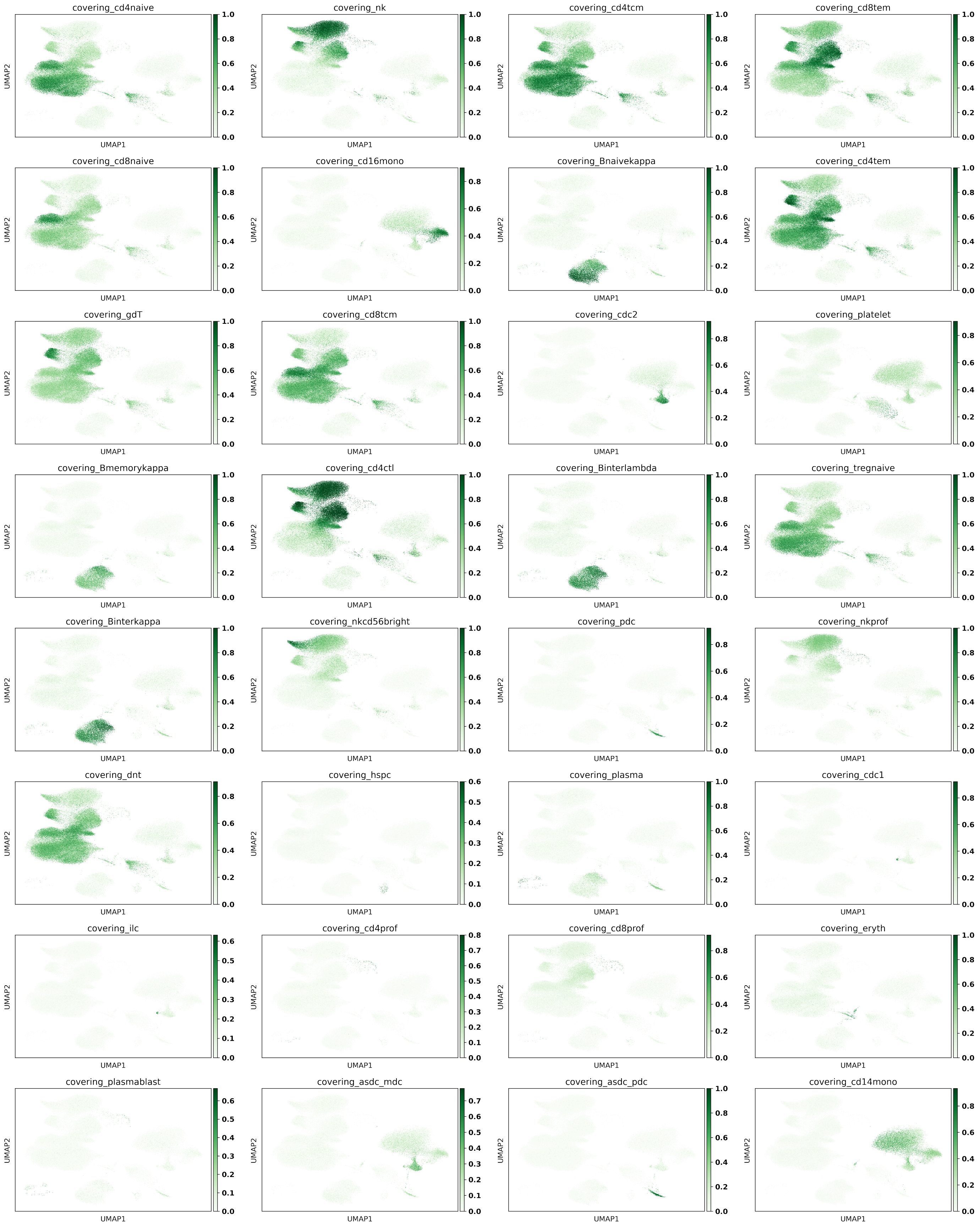


**Supplementary Figure 21: Transfer of CellCover marker panels from PBMC CITE-seq dataset to Stephenson COVID-19 PBMC dataset.** For each of the 32 cell types in the PBMC datasets, a CellCover marker panel of depth 5 and covering rate of 98% was derived. Each panel was transferred to the target dataset, with the plots showing the proportion of marker genes expressed in cells of the target dataset, as indicated by the titles of each subplot.

### 10 oRG vs Gliogenic Across Mammalian Species

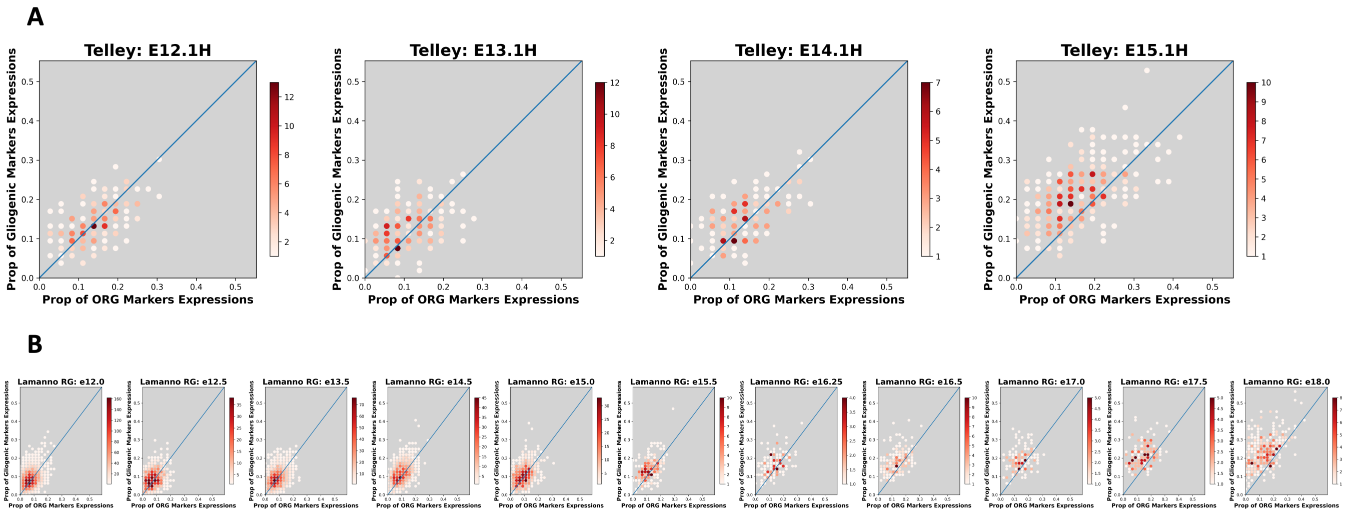


**Supplementary Figure 22**: **Transfer of oRG and gliogenic marker panel, derived from the sorted cell types of human fetal cortex, to mouse progenitor cells.** The intensity of the dot at coordinate (x,y) indicates the number of mouse progenitor cells with $(100 \times x)$ percent of oRG markers expressed and $(100 \times y)$ percent of gliogenic markers expressed. A) Transfer to the four embryonic dates of 1H cells in Telley dataset. B) Transfer to the LaManno radial glia cells across time.

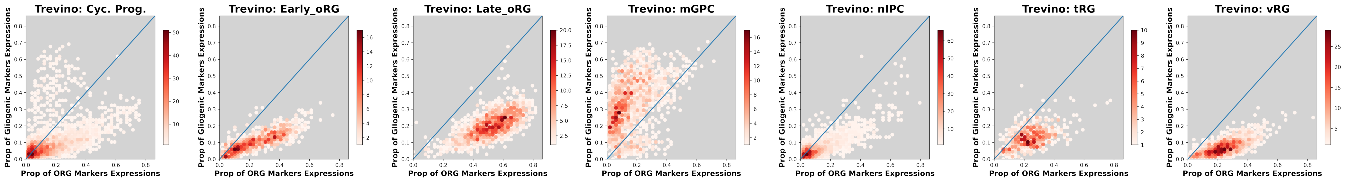


**Supplementary Figure 23**: **Transfer of oRG and gliogenic marker panel, derived from the sorted cell types of human fetal cortex, to human progenitor cell subtypes.** The intensity of the dot at coordinate (x,y) indicates the number of human progenitor cells with $(100 \times x)$ percent of oRG markers expressed and $(100 \times y)$ percent of gliogenic markers expressed.

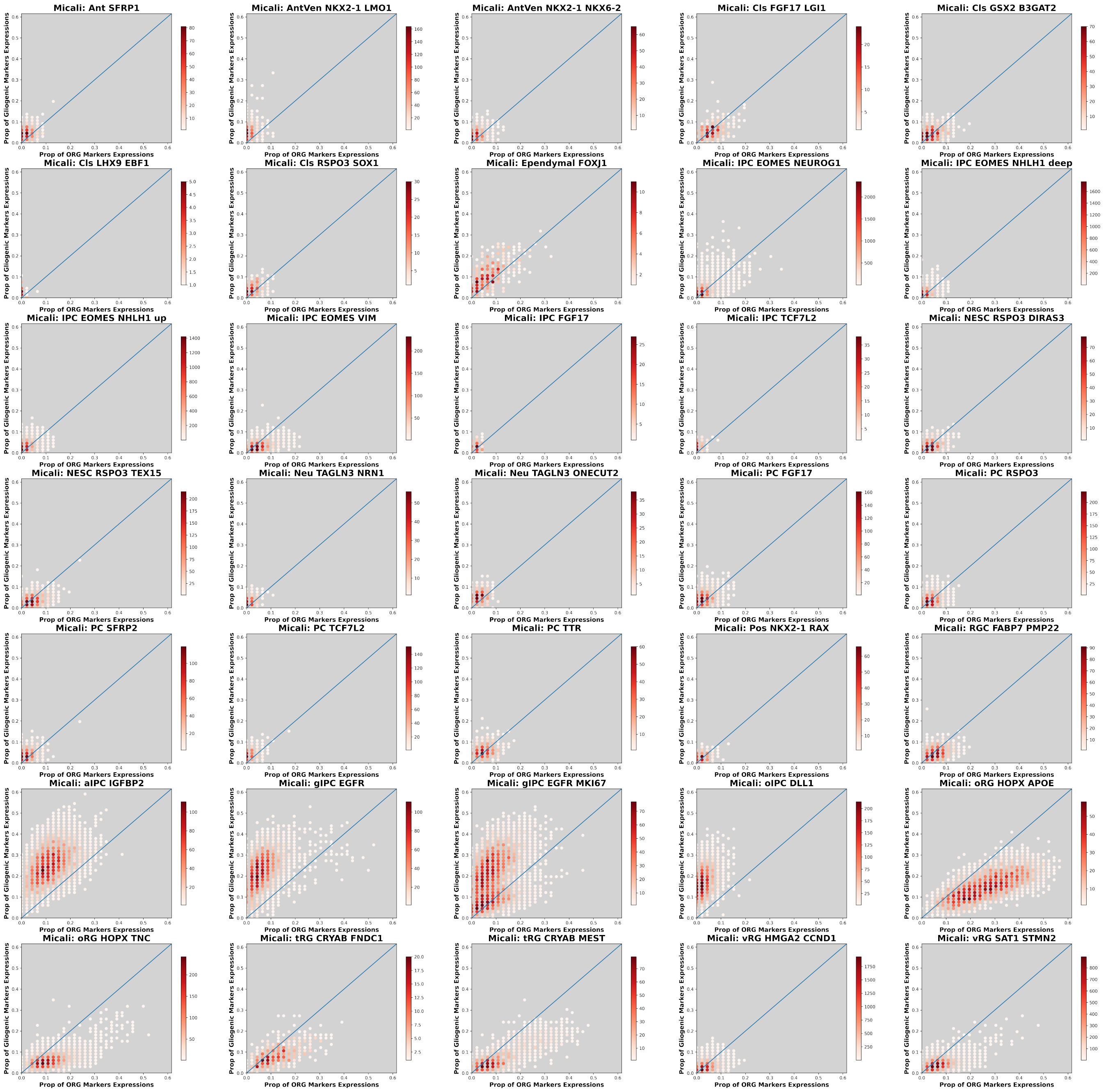


**Supplementary Figure 24: Transfer of oRG and gliogenic marker panel, derived from the sorted cell types of human fetal cortex, to macaque progenitor cell subtypes.** The intensity of the dot at coordinate (x,y) indicates the number of macaque progenitor cells with $(100 \times x)$ percent of oRG markers expressed and $(100 \times y)$ percent of gliogenic markers expressed.
